# Identifying digenic disease genes using machine learning in the undiagnosed diseases network

**DOI:** 10.1101/2020.05.31.125716

**Authors:** Souhrid Mukherjee, Joy D Cogan, John H Newman, John A Phillips, Rizwan Hamid, Undiagnosed Diseases Network, Jens Meiler, John A. Capra

## Abstract

Rare diseases affect hundreds of millions of people worldwide, and diagnosing their genetic causes is challenging. The Undiagnosed Diseases Network (UDN) was formed in 2014 to identify and treat novel rare genetic diseases, and despite many successes, more than half of UDN patients remain undiagnosed. The central hypothesis of this work is that many unsolved rare genetic disorders are caused by multiple variants in more than one gene. However, given the large number of variants in each individual genome, experimentally evaluating even just pairs of variants for potential to cause disease is currently infeasible. To address this challenge, we developed DiGePred, a random forest classifier for identifying candidate digenic disease gene pairs using features derived from biological networks, genomics, evolutionary history, and functional annotations. We trained the DiGePred classifier using DIDA, the largest available database of known digenic disease causing gene pairs, and several sets of non-digenic gene pairs, including variant pairs derived from unaffected relatives of UDN patients. DiGePred achieved high precision and recall in cross-validation and on a held out test set (PR area under the curve >77%), and we further demonstrate its utility using novel digenic pairs from the recent literature. In contrast to other approaches, DiGePred also appropriately controls the number of false positives when applied in realistic clinical settings like the UDN. Finally, to facilitate the rapid screening of variant gene pairs for digenic disease potential, we freely provide the predictions of DiGePred on all human gene pairs. Our work facilitates the discovery of genetic causes for rare non-monogenic diseases by providing a means to rapidly evaluate variant gene pairs for the potential to cause digenic disease.

## INTRODUCTION

Causal genetic variants have been identified for thousands of Mendelian diseases (Ionita-Laza et al., 2011; Ng et al., 2009, 2010). However, in spite of the advent of cheaper and more accurate sequencing technologies, causal variants have not been identified for approximately half (~3000) of known rare genetic diseases (Boycott et al., 2017, 2019; Chong et al., 2015). To help address this challenge, the Undiagnosed Diseases Network (UDN) was established by the NIH in 2014. Comprising teams of researchers and clinicians from 12 sites across the United States, the UDN integrates whole exome/genome sequencing with expert clinical evaluation to develop diagnoses and treatment plans for patients, who could not be diagnosed by conventional clinical approaches (Gahl et al., 2015, 2016; Ramoni et al., 2017). Although this approach has yielded much success, (Bostwick et al., 2017; Chao et al., 2017; Johnston et al., 2018; Küry et al., 2017; Liu et al., 2018; Machol et al., 2018; Marcogliese et al., 2018; Oláhová et al., 2018b, 2018a; Poli et al., 2018; Pomerantz et al., 2018; Schoch et al., 2017; Tokita et al., 2018) more than half of all UDN cases remain undiagnosed. We hypothesize that many of these unsolved, rare cases might involve variants in multiple genes that only when combined result in a disease phenotype complicating diagnosis.

Variants in multiple genes can synergistically lead to disease via many different mechanisms (Auer et al., 2018; Badano and Katsanis, 2002; van Heyningen and Yeyati, 2004; Pehlivan et al., 2019). Digenic inheritance was first observed in 1994, when concurrent mutations in two genes were found to be responsible for causing retinitis pigmentosa.(Kajiwara et al., 1994) Digenic inheritance is the simplest form of oligogenic inheritance in which variants in multiple genes lead to disease.(Gazzo et al., 2016; Lupski, 2012; Schäffer, 2013) There are various classifications of digenic disease,(Deltas, 2018) but in all cases of digenic inheritance the phenotype results from variants in two genes. In isolation, the individual variants that form a digenic pair are benign or lead to a simpler phenotype. However, upon simultaneous mutation, the variants either interact to produce disease or combine to produce a more complex, and usually more severe, phenotype that cannot be explained by variants in one gene alone.

The Digenic Diseases Database (DIDA) (Gazzo et al., 2016) has chronicled several hundred cases of digenic disease. Analyses of DIDA have revealed that digenic disease causing gene pairs are more likely to functionally and/or physically interact with one another than expected by chance (Gazzo et al., 2016). Machine learning approaches have been developed to distinguish between different types of digenic disease pairs (Gazzo et al., 2017) and to identify disease causing variant combinations, (Boudellioua et al., 2018; Papadimitriou et al., 2019) including oligogenic combinations of greater than two genes (Renaux et al., 2019).

We hypothesize that the disease phenotype in some unresolved UDN patients is likely a result of digenic inheritance and develop DiGePred, a high-throughput machine learning tool for evaluating the likelihood that dysfunction of gene pairs leads to digenic disease. We focus on the specific challenge of identifying gene pairs that have functional or phenotypic synergy to cause a digenic disease when both are disrupted in a patient. Our approach is based on supervised machine learning using a random forest classifier trained on diverse functional, network, and evolutionary properties of known digenic gene pairs versus realistic sets of non-digenic gene pairs, including variant pairs from healthy individuals. We demonstrate that DiGePred is accurate and that, in comparison to recent approaches, it has a low false positive rate, which is essential for clinical applications. To aid in rapid screening of patients for potential digenic disease variants, we provide a classification of the digenic disease potential for all human gene pairs.

## RESULTS

### Digenic disease gene pairs have different attributes than non-digenic disease gene pairs

Our goal in this study is to develop a machine learning classifier for identifying gene pairs that are likely to cause disease when both are disrupted simultaneously, but fail to produce a strong disease phenotype when disrupted in isolation. To this end, we consider all unique known digenic disease pairs curated by the DIDA database and contrast them with several sets of non-digenic disease pairs. Since our ultimate application is the detection of potential digenic diseases in patients, most of our results focus on comparisons of known digenic gene pairs and gene pairs with variants in 58 “unaffected” parents, siblings, and other relatives of 25 UDN patients (**Figure 1A**). However, as we show below, our results are similar using other strategies for defining non-digenic disease gene pairs.

**FIGURE 1.**
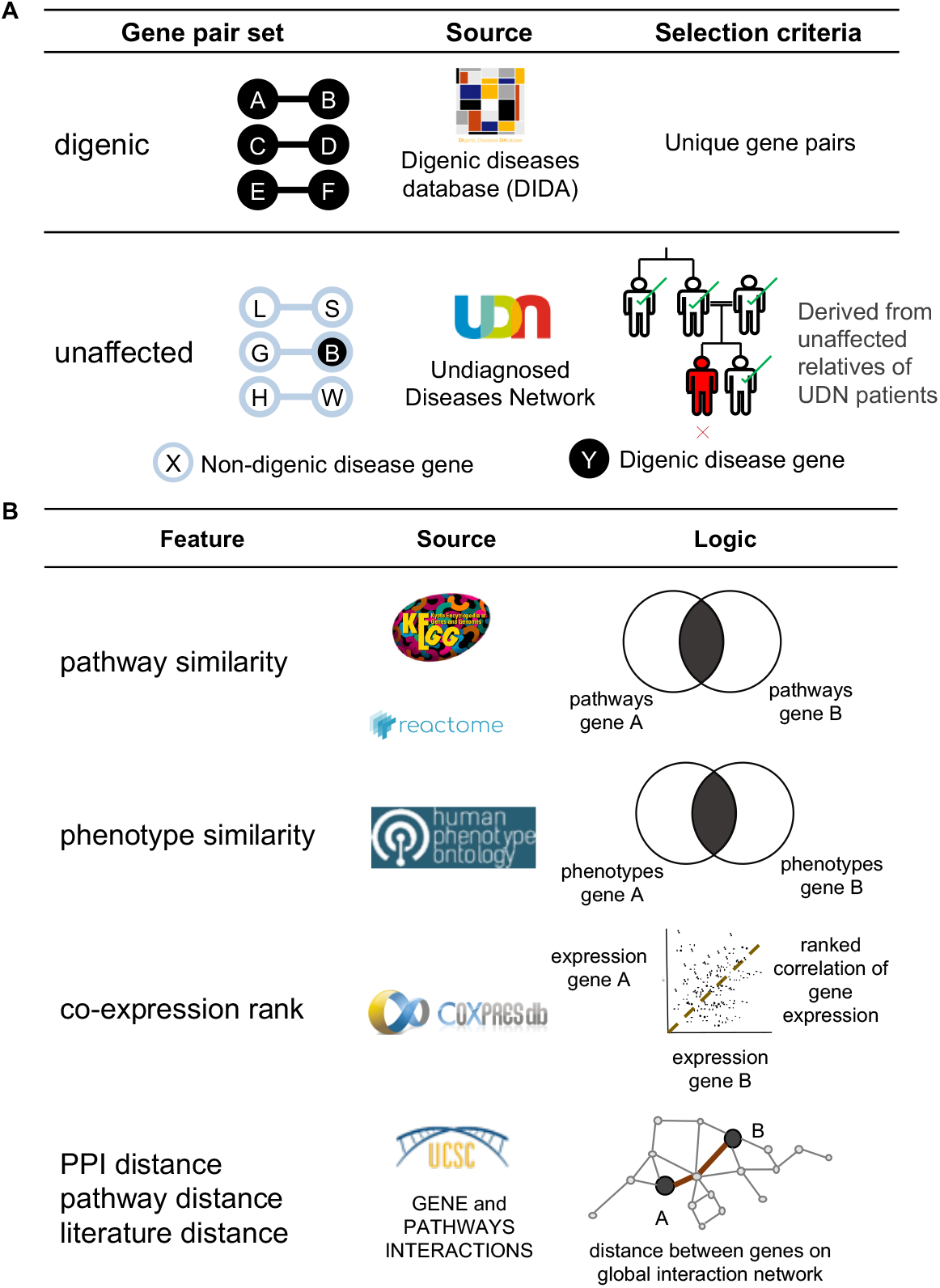
Training sets and features used for random-forest-based identification of digenic disease gene pairs. **(A)**. The digenic gene pairs (positives) were derived from the Digenic Diseases Database (DIDA). Unique gene pair combinations (n=140) were used for training and testing. The likely non-digenic gene pairs (negatives) were derived from unaffected relatives of UDN patients. Genes with rare variants in the same individual were used as an unaffected non-digenic gene pair. We also considered several others of negative training examples (**Figure S2**). **(B)**. We considered six network and functional features (NFFs) for training the first digenic disease classifier: i) *pathway similarity*: Jaccard similarity of pathway annotations from KEGG and Reactome for both genes; ii) *phenotype similarity*: Jaccard similarity of phenotype annotations from HPO for both genes, iii) *co-expression rank*: co-expression rank of gene pair compared to all other gene pairs across multiple tissues from COXPRESdb; iv-vi) network distances between the genes on protein-protein, pathway, and literature mined interaction networks from UCSC gene and pathway interaction browser database. Subsequent classifiers considered additional evolutionary and functional features (**Figure S3**).

Pairs of genes harboring mutations known to cause digenic disease have distinct biological properties when compared with random gene pairs (Gazzo et al., 2016). Previous work has shown that digenic disease pairs have high protein interaction network connectivity and proximity. More than 35% of known digenic disease pairs directly interact on a protein-protein interaction (PPI) network, and ~60% of digenic gene pairs are one gene away on the interaction network. Similarly, ~20% of digenic pairs are in the same biochemical pathway, and ~40% are expressed in the same tissues (Gazzo et al., 2016).

Based on this prior knowledge we devised a list of six “network and functional features” (NFFs) to use as attributes for distinguishing between digenic and non-digenic gene pairs (**Figure 1B**): 1) *Pathway Similarity*, defined as the Jaccard similarity (Jaccard, 1912) between the genes’ membership in ~1800 pathways from KEGG (Kanehisa et al., 2017) and Reactome (Fabregat et al., 2018; Milacic et al., 2012); 2) *Phenotype Similarity*, the Jaccard similarity between the ~6000 phenotypes from Human Phenotype Ontology (HPO) (Köhler et al., 2017) associated with the genes; 3) *Co-expression Rank*, defined as the rank of the co-expression of the genes across 23 co-expression platforms compared to other gene pairs from COXPRESdb (Okamura et al., 2015); 4) *PPI Distance*, the distance on a global PPI network; 5) *Pathway Distance*, the distance on an annotated biochemical pathway network; and 6) *Literature Distance*, the distance on a literature-mined interaction network, derived from the UCSC gene and pathway interaction database (Poon et al., 2014).

We compared the distribution of the NFFs for digenic and non-digenic gene pairs from unaffected relatives of UDN patients, and as expected from previous work, the distribution of each NFF was significantly different between digenic and non-digenic pairs (**Figure S1**; *P* < 10^−20^ for each, Mann-Whitney U (MWU) test). This suggests that a machine learning approach may enable distinguishing digenic from non-digenic disease pairs.

### Random forest classifiers accurately identify digenic pairs using network and functional features

We trained the random forest machine learning classifier using the six NFFs to distinguish 140 digenic disease gene pairs (positives) from ~8,400 non-digenic gene pairs, derived from genes with rare variants (allele frequency <1%) in unaffected relatives of UDN patients (negatives). The large class imbalance (~1:75) reflects the expectation that only a very small fraction of gene pairs are likely to produce digenic disease. For example, comprehensive studies of genetic interactions have found that one in approximately 40 gene pairs interact.(Costanzo et al., 2016)

We divided the available gene pairs into training (64%), validation (16%), and testing sets (20%). We trained, evaluated, and compared different models using 10-fold cross-validation within the training and validation sets (**Figure 2**). The testing set was only analyzed after models had been finalized. We evaluated performance using Receiver Operating Characteristic (ROC) and Precision-Recall (PR) curves.

**FIGURE 2.**
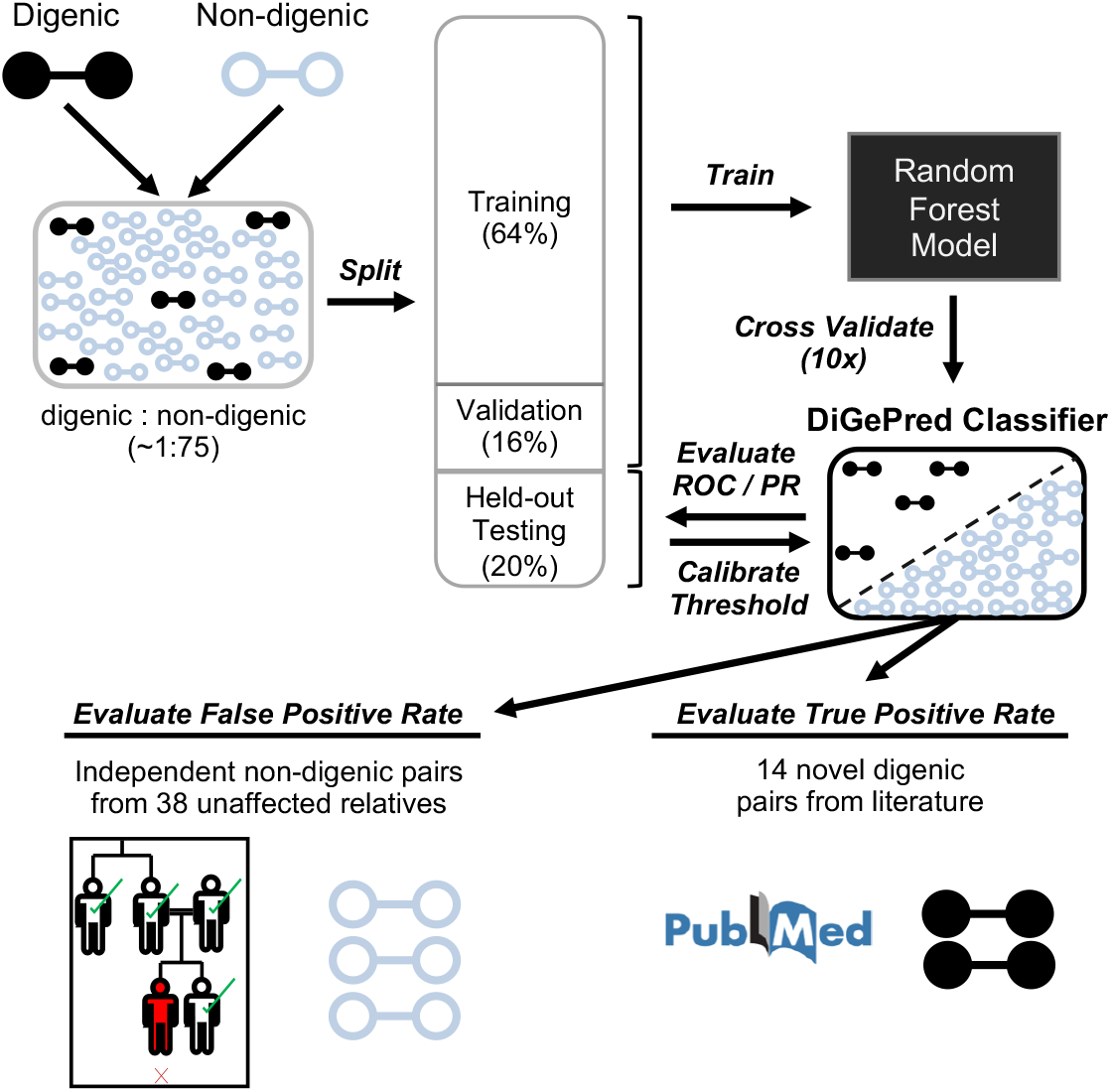
Schematic of the protocol for training and evaluating the DiGePred digenic disease pair classifier. Known digenic pairs (positives) and variant gene pairs from healthy individuals (negatives) were combined at ~1:75 ratio. The combined pairs were divided into training (64%), validation (16%) and held-out testing datasets (20%). The DiGePred random forest classifier was trained and cross-validated using the training and validation sets. The final performance estimate for the trained DiGePred classifier was quantified by the area under the Receiver Operator Characteristic (ROC) and Precision-Recall (PR) curves (AUCs) on the held-out test set. This set was also used to establish suggested thresholds on the continuous DiGePred score. DiGePred’s potential clinical utility was further demonstrated by applying it to an additional positive set of 14 novel digenic pairs from the recent literature and an external non-digenic set of gene pairs from 38 unaffected relatives of UDN patients.

The random forest classifier distinguished digenic and non-digenic gene pairs very accurately using the six NFFs. It achieved an average ROC area under the curve (AUC) of 0.91 and a PR AUC of 0.69 on average over 10 folds of cross-validation (**Figure 3**). Though the PR AUC is lower than the ROC AUC, the algorithm retains near perfect precision at recall above 60% (**Figure 3B**).

**FIGURE 3.**
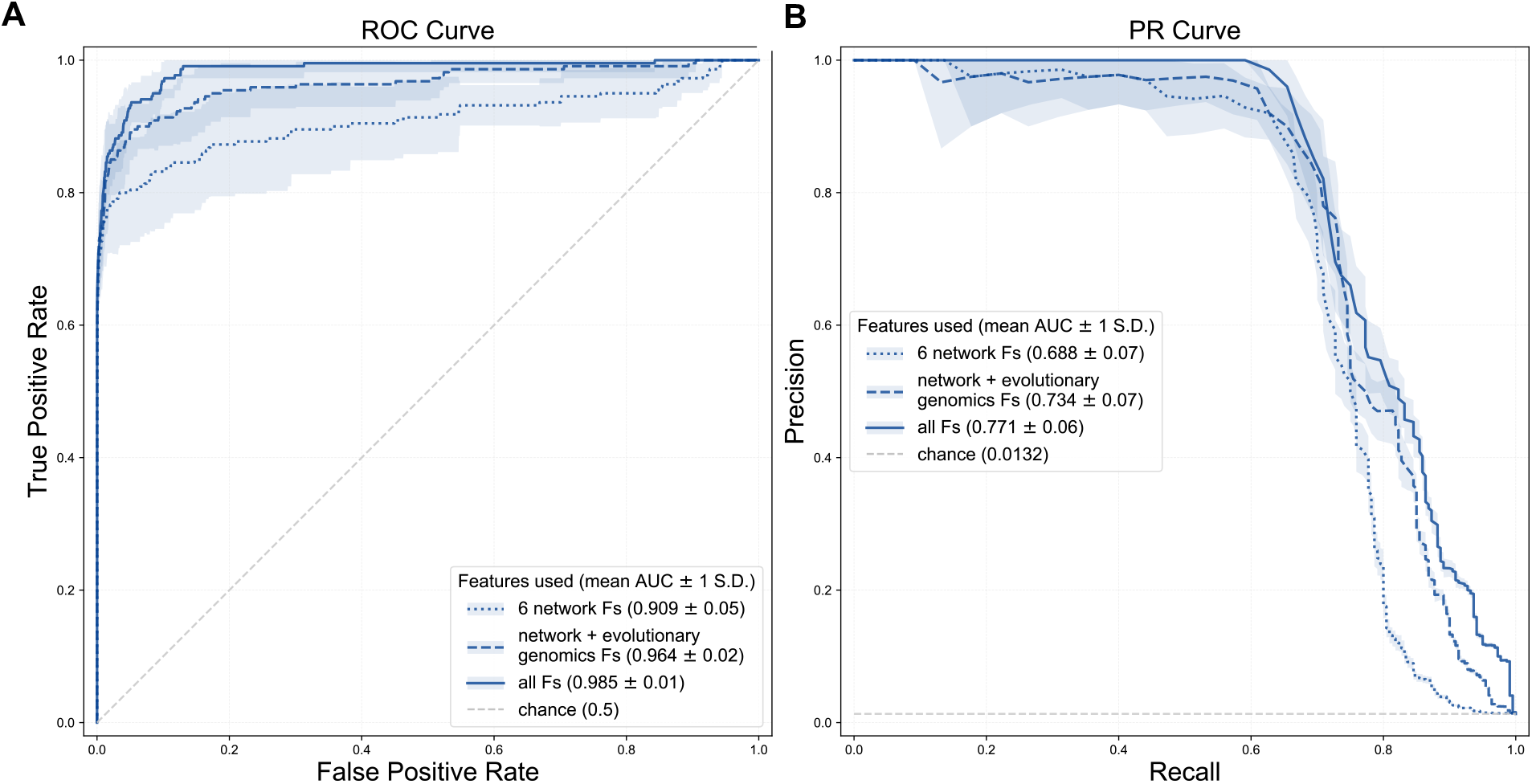
Random forest classifiers can accurately distinguish digenic and non-digenic gene pairs using different feature sets. Performance of the classifier at distinguishing between known digenic pairs and gene pairs from healthy individuals trained using different feature sets as evaluated by: **(A)** Receiver Operating Characteristic (ROC) curves and **(B)** Precision-Recall (PR) curves. Classifiers trained on three sets of features are compared: i) six network and functional features (NFFs) (dotted line); ii) the six NFFs and evolutionary genomics features; and iii) the six NFFs, evolutionary genomics features, and gene-level network and functional features. The mean curves across 10 cross-validation folds are plotted with shaded areas representing the standard deviation.

### Including additional features improves ability to identify digenic disease genes

The performance of the classifier based on the six NFFs alone was strong; however, there are many other sources of biological information beyond the NFFs that could potentially inform either the nature of the relationship between genes or the relative likelihood and risk of a gene being mutated and causing disease. We tested if additional features in training the classifier would increase performance and contribute to the robustness of the classifier.

First, we trained classifiers using the six NFFs and five additional evolutionary features that reflect the evolutionary history and constraint on the genes (**Figure S3**). These features were: 1) the evolutionary ages of the genes; 2) their essentiality; 3) their intolerance to loss of function mutations, 4) the selection pressure acting on them through mammalian evolution (dN/dS) and 5) their haploinsufficiency scores. Since each of these features applies to a single gene (rather than a pair), we created two features for each pair. The addition of evolutionary features substantially improved classifier performance: average ROC AUC of 0.96 and PR AUC of 0.73 (**Figure 3**).

Next, we considered additional features derived from network and functional annotations of the gene pairs (**Figure S3**). These features were designed to add additional gene-focused (rather than gene-pair-focused) context and explore the sufficiency of the six NFFs. These features were: 1) the number of pathways, 2) phenotypes, 3) network neighbors, and 4) genes co-expressed for each individual gene in a candidate pair. Considering these features also further improved classifier performance, with an average ROC AUC of 0.99 and PR AUC of 0.77 with all features (**Figure 3**).

### Digenic disease genes can be distinguished from many non-digenic gene sets

To further explore the properties of digenic disease genes and the ability of our classification approach to recognize them, we defined three additional sets of non-digenic disease gene pairs (**Figure S2**). First, we created a “random” set of non-digenic gene pairs by randomly selecting gene pairs from all possible human genes (excluding known digenic pairs). Second, we constructed a “permuted” non-digenic set by generating all possible gene pairs from genes known to be involved in a digenic gene pair, and removing the pairs known to be digenic. Third, we created a “matched” non-digenic gene pair set that closely matched the NFF distributions of the digenic gene pairs (**Figure S4**). However, we note that we were not able to match the distribution of all NFFs perfectly given the skewed distribution of the digenic disease pairs. The matched set enables exploration of how well our classification approach can identify digenic pairs among non-digenic pairs with similar NFF distributions.

We used the same training and evaluation approach as described for the unaffected negative set to train random forest classifiers to distinguish digenic disease gene pairs from each of these three additional negative sets using all the network, functional, and evolutionary features. In each case, the classifiers performed very well (**Figures 4**, **S5**). As expected, the classifier trained to distinguish digenic pairs from random pairs performed the best (average ROC AUC of 0.98 and PR AUC of 0.73). Given the similar attributes between the digenic disease and the other negative sets, the permuted and matched classifiers performed slightly less well, but still achieved very strong performance with average ROC AUCs of 0.98 and 0.98 and PR AUCs of 0.59 and 0.61, respectively.

**FIGURE 4.**
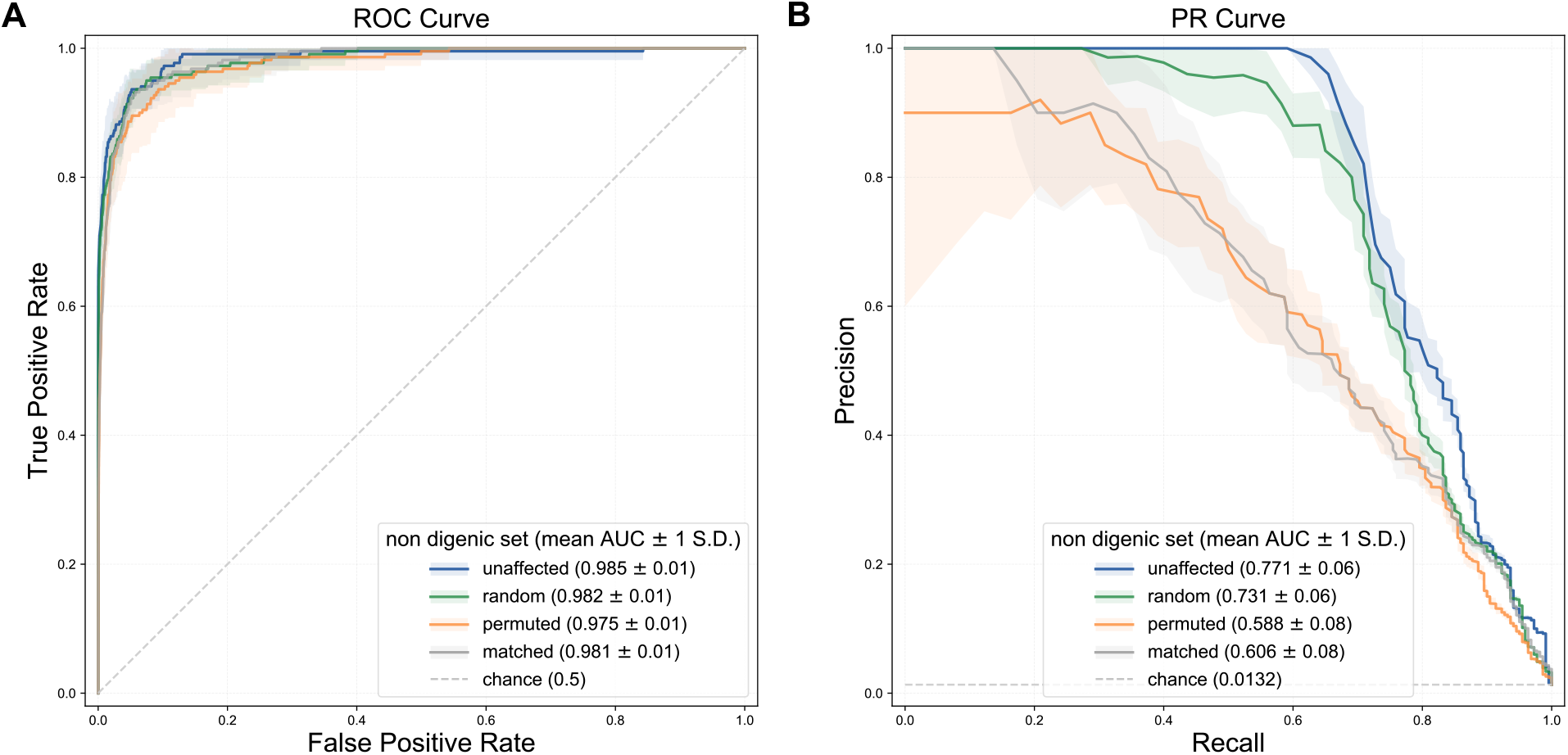
Digenic disease gene pairs can be distinguished from different non-digenic sets. Performance of random forest classifiers at distinguishing between digenic pairs and different non-digenic sets trained using the full set of features as evaluated by **(A)** ROC and **(B)** PR curves. We considered four different negative sets: i) *Unaffected*, derived from healthy relatives of UDN patients (blue); ii) *Random*, derived by randomly selecting pairs of genes (green); iii) *Permuted*, derived by generating permutations of known digenic pairs (orange); iv) *Matched*, derived by matching the distribution of network and functional features observed among the digenic pairs (grey). The mean curves across 10 cross-validation folds are plotted with shaded areas representing the standard deviation. The Permuted and Matched negative sets provide a greater challenge given their similarities with the digenic pairs, but the approach still achieved strong performance at these tasks. The feature importance values highlight phenotypic and pathway similarity and are similar for each classifier, except the Matched classifier (**Figure S6**).

### Feature importance varies for classifiers trained on different non-digenic sets

We estimated the importance of the features to the classifiers using the mean decrease in node impurity approach (**Figure S6**). For the classifier trained using variant gene pairs from unaffected relatives, the number of phenotypes shared between the pair, number of pathways shared, and the overall number of annotated phenotypes were the most important features (28%, 9%, and 9% of the weight, respectively). The feature importance values were similar for the classifiers trained using random gene pairs and permuted digenic gene pairs (**Figure S6**).

The feature importance values were most different for the matched classifier with significantly lower feature importance for the NFFs. This was expected, because the differences between the positive and negative training examples in individual NFFs were minimal for this classifier. Instead, a range of evolutionary and individual gene-level functional features took on similar levels of importance (**Figure S6**). This indicates that information in gene-level features related to evolution, gene importance, and relevance to physiology contain useful information about the likelihood of gene pairs interacting to produce digenic disease.

### DiGePred accurately identifies held-out digenic pairs

Based on the previous results, the best performing model was the classifier trained on the unaffected non-digenic gene pairs using all the features. Furthermore, this classifier most closely reflects the distribution of gene mutations likely to be seen in real clinical applications. To obtain an unbiased estimate of its performance, we evaluated it using held-out sets of digenic and non-digenic pairs. These sets were not used for training or validating the classifier and maintained the ~1:75 ratio used during training. Since we have a large number of unaffected gene pairs, we repeated the evaluation 100 times using the same set of digenic pairs (n=28), but unique sets of unaffected non-digenic gene pairs.

The mean ROC AUC for the evaluation on the held-out sets was 0.994, while the mean PR AUC was 0.91 (**Figure 5**). We evaluated the performance of classifiers trained on the other non-digenic gene pair sets on their corresponding held-out sets, and the ROC AUCs were > 0.97 and PR AUCs were > 0.50 in all cases. (**Figure S7**) We also trained and evaluated classifiers on datasets constructed so that no individual genes overlapped between the training and testing sets. As expected, this task was more difficult, but performance was still very strong with ROC AUC of 0.96 and PR AUC of 0.68 (**Figure S8**).

**FIGURE 5.**
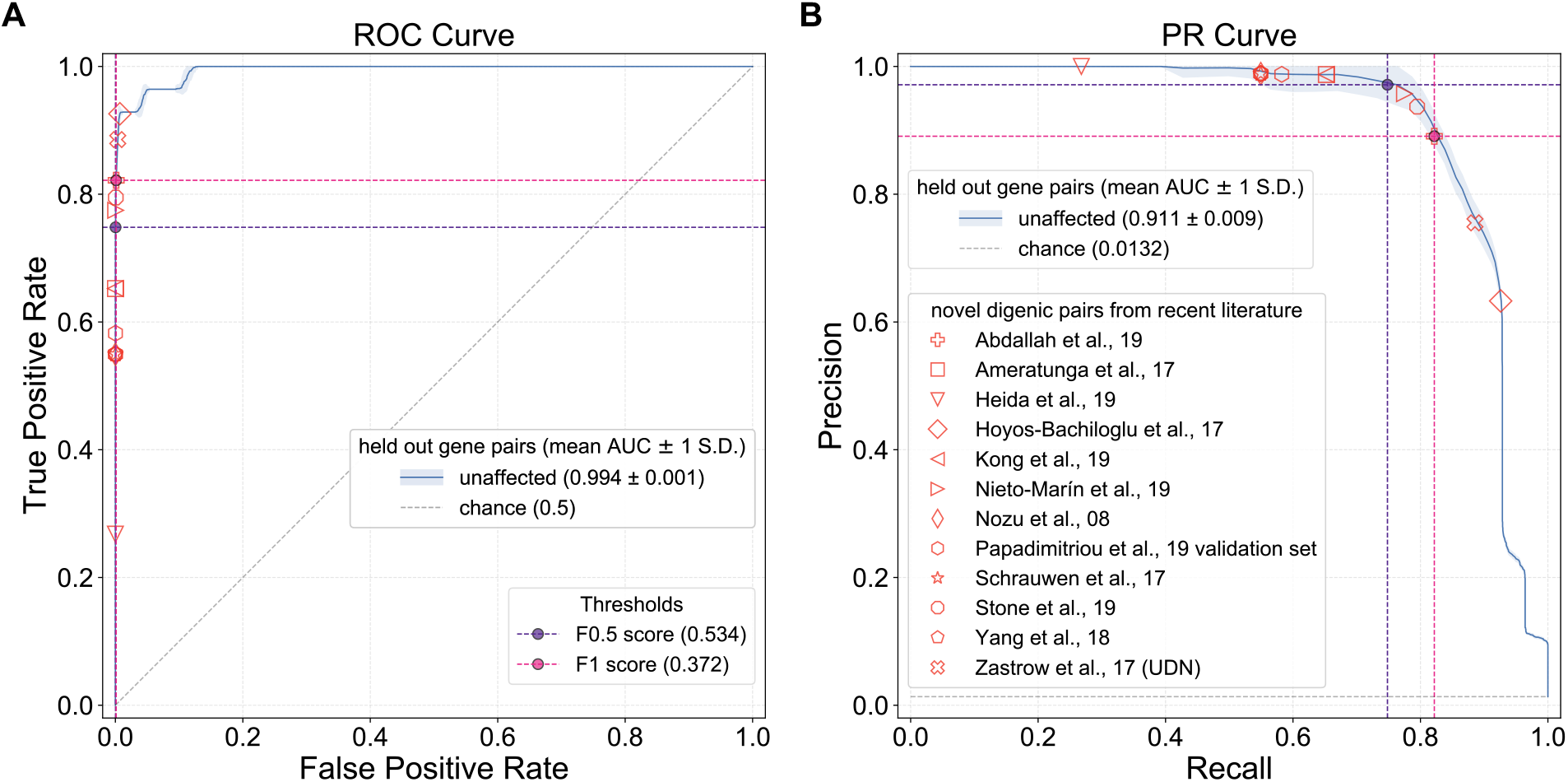
DiGePred performs well on the held-out test set and accurately identifies novel digenic pairs from the recent literature. Performance of DiGePred on the held-out test set as evaluated by **(A)** ROC and **(B)** PR curves. The DiGePred classifier was trained using all features and the unaffected set as negatives. Geometric shapes with red borders indicate the position on the ROC and PR curves corresponding to the DiGePred scores assigned to 13 novel digenic pairs reported in the recent literature. The pink and purple circles represent the points on the curves corresponding to the DiGePred score thresholds that maximize the F_1_ (0.372) and the F_0.5_ (0.534) metrics. Given the importance of precision in clinical applications, we propose the score maximizing the F_0.5_ metric or higher as a threshold for calling a gene pair digenic. At this threshold 9 of the 13 novel digenic pairs are predicted to be digenic with a low expected false positive rate (<=0.012%). All but one digenic pair score above the F_1_ threshold. The Zastrow et al. 2017 pair (red x) is not truly digenic, but represents a patient with pathogenic variants in two genes that do not interact.

To establish prediction thresholds on the output of this classifier, we computed thresholds that maximize the F_1_ and F_0.5_ scores. The F_1_ is maximized at a digenic score of 0.369, and the F_0.5_ is maximized at a digenic score of 0.534. Since we anticipate that precision is more important than recall in most applications, we suggest use of the F_0.5_-based threshold. At this threshold, the classifier correctly identified 20 of 28 digenic gene pairs in the held-out test set, with a false positive rate of 0.026% (**Figure 5, Supplementary Table 1**). We refer to this model as the DiGenic Predictor (DiGePred).

### DiGePred identifies novel digenic pairs from the recent literature

While the test set was not seen by the classifier prior to evaluation, it was still obtained from DIDA, the source of digenic pairs for training and testing. Thus, we further applied our classifier to 13 new digenic pairs obtained from recent literature, not included in DIDA (**Supplementary Table 2**). We derived three digenic pairs (*(CEP290, RPE65), (AHI1,CEP290), (CEP290, CRB1)*) from the validation set used by a recently published digenic classifier (Papadimitriou et al., 2019). The other digenic gene pairs (*(CLCNKA, CLCNKB), (TCF3, TNFRSF13B), (IFNAR1, IFNGR2), (PCDH15, USH1G), (LAMA4, MYH7), (KCNE2, KCNH2), (CLCNKB, SLC12A3), (CACNA1C, SCN5A), (FGFR1, KLB), (CLCN7, TCIRG1)*) were derived from recently reported cases of digenic disease, respectively: (Abdallah et al., 2019; Ameratunga et al., 2017; Heida et al., 2019; Hoyos-Bachiloglu et al., 2017; Kong et al., 2019; Nieto-Marín et al., 2019; Nozu et al., 2008; Schrauwen et al., 2018; Stone et al., 2019; Yang et al., 2018).

DiGePred correctly identified 9 of the 13 novel digenic pairs at the F_0.5_ threshold, with an expected false positive rate of 0.012% or lower. (**Figure 5**). Two of the gene pairs missed at the F_0.5_ threshold, *CACNA1C* and *SCN5A* (Nieto-Marín) and *FGFR1* and *KLB* (Stone) were identified as digenic at the F_1_ threshold, at FPRs of 0.034% and 0.056%, respectively. Among the two other pairs missed, IFNAR1 and IFNGR2 (Hoyos-Bachiloglu) had very low phenotype coverage, while *LAMA4* and *MYH7* (Abdallah) was identified as digenic by the matched classifier (**Figure S9**).

While searching the recent literature for novel digenic pairs, we also found a case that did not meet the strict criteria for digenic interaction, but that did have fucntional synergy between two genes leading to a disease phenotype. The gene pair comes from a solved UDN case in which variants in *FBN1* and *TRPS1* were implicated in the patient phenotype, but these variants did not interact (Zastrow et al., 2017). DiGePred identified this gene pair as non-digenic; nonetheless, it had a higher digenic score than 99% of the non-digenic gene pairs (**Supplementary Table 3**), suggesting the potential of the classifier to highlight pairs of functionally related genes.

### DiGePred has a low false positive rate in real-world applications

Individuals often carry hundreds of protein-coding variants of unknown significance, which results in thousands of potential digenic disease pairs per individual. Thus, when considering the application of classifiers to individuals’ genomes, it is essential to understand and control the false positive rate. To this end, we evaluated DiGePred on gene pairs with rare variants predicted to disrupt protein function in 38 human genomes from unaffected parents and relatives of UDN patients not used in training the algorithm. These healthy individuals should not contain any true digenic disease pairs, so any positive predictions on gene pairs from these individuals are very likely to be false positives. The gene pairs from these individuals were not used in the training, testing, or held-out sets.

At the F_0.5_ threshold, 13% of unaffected individuals had no predicted candidate digenic pairs and 50% had only one candidate digenic pair. On average, two digenic pairs were predicted per individual, and only two individuals had more than five digenic pairs (**Figures 6, S10**). Furthermore, no gene pairs with rare variants have scores above 0.9 (**Figure S10**). This suggests that users can adjust the score threshold to reflect their tolerance for false positives. In contrast, we applied the recently published ORVAL (Papadimitriou et al., 2019; Renaux et al., 2019) method for identifying digenic disease pairs to variants from these same individuals. At its highest confidence threshold, ORVAL predicted that all these healthy individuals have digenic disease pairs, with an average of 855 highest confidence digenic pairs per individual and more than a thousand digenic pairs predicted for 28 (~74%) individuals, with all of the individuals being predicted with > 300 digenic pairs. (**Figure 6**). This is a significantly larger number of candidate digenic disease pairs per individual than DiGePred (*P* = 2.07×10^−14^, MWU test), and these are all very likely to be false positives. This difference in number of false positives was recapitulated for all gene selection criteria and all models of training considered (**Figures S11-16**).

**FIGURE 6.**
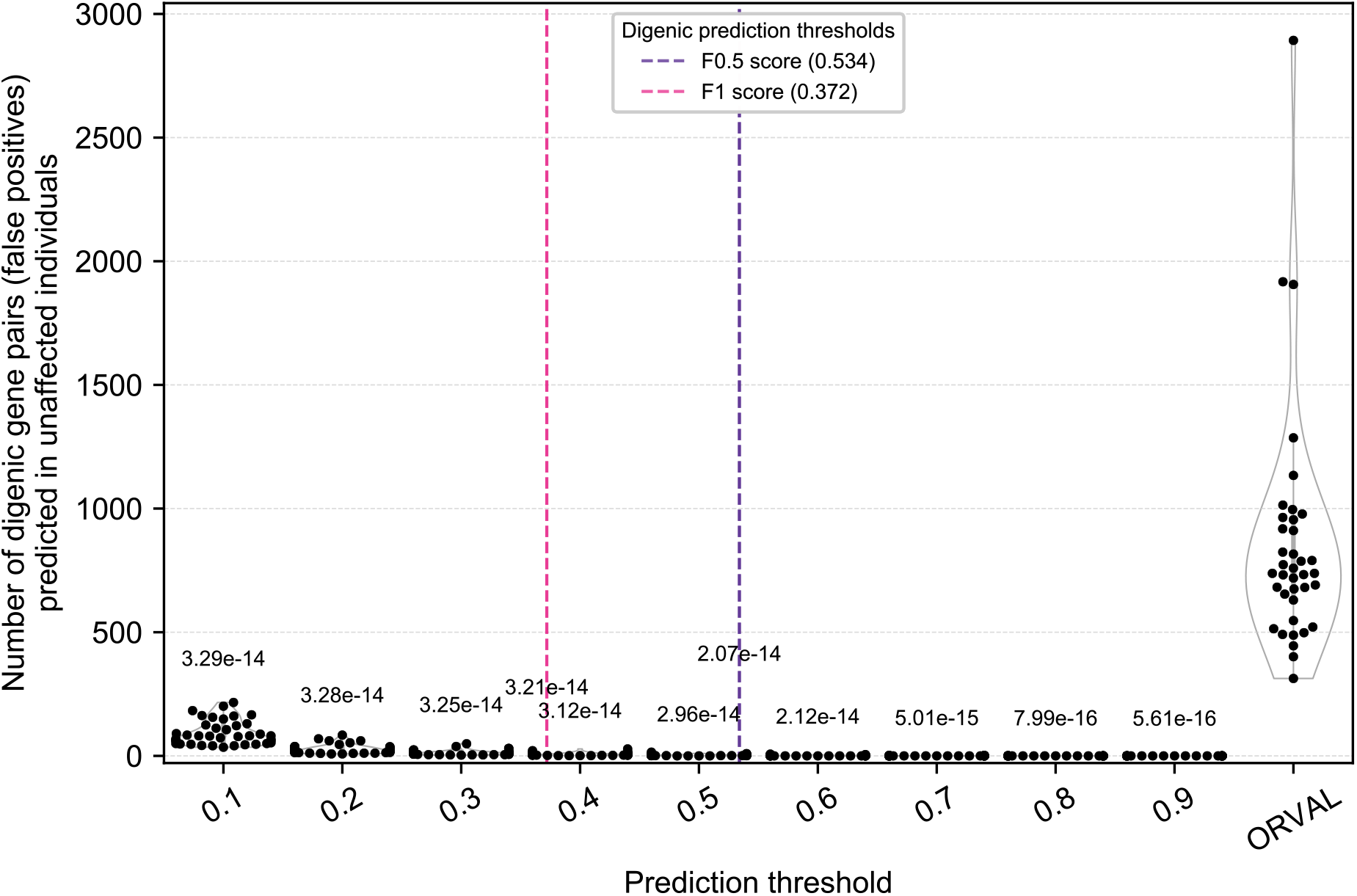
DiGePred has a low false positive rate and outperforms a recent digenic gene prediction method. The number of digenic pairs identified for each of 38 healthy relatives of UDN patients is plotted at a range of DiGePred thresholds (x-axis) and for the ORVAL/VarCOPP method. The score thresholds that maximize the F_1_ and F_0.5_ metrics on the held out data are shown in pink and purple, respectively. Since these individuals are healthy, any predicted digenic disease pairs are very likely false positives. DiGePred predicts significantly fewer digenic pairs at each threshold than ORVAL (Mann-Whitney U test, p-values above each bar). At the F_0.5_ threshold, DiGePred predicts an average of two digenic pairs per healthy individual and none above the 0.9 threshold, while ORVAL predicts an average of 855 digenic pairs per healthy individual at its strictest threshold (**Figure S10**). Results were similar for classifiers trained on other negative sets (**Figures S11-16**).

### Prediction of digenic pairs among all human gene pairs at various confidence thresholds

To aid in the rapid evaluation of digenic disease potential for a pair of genes of interest, we updated DiGePred by training the method using all digenic pairs from DIDA (to make use of all known digenic pairs) and variant gene pairs from healthy relatives of UDN patients. We applied DiGePred to all possible human gene pairs. A gene pair was deemed a candidate digenic pair if the digenic score met the F_0.5_ threshold as described above. As expected, the percentage of all possible gene pairs that were identified as digenic at our most confident threshold was very low (33,272 out of 155.33 million gene pairs, 0.021%). These predictions and the raw digenic scores are available in **Supplementary Table 4.**

Overall, 4336 genes are involved in at least one predicted digenic pair. This illustrates that DiGePred is not just prioritizing gene pairs that include a gene in a known digenic pair. In fact, only 8 of the top 100 genes with the most predicted digenic pairs occur in DIDA. These genes are enriched for Gene Ontology functional annotations “translational initiation” and “structural component of ribosome”. For example, the gene *ARID1B*, which has the highest number of predicted digenic pairs, with 188, encodes a component of the SWI/SNF chromatin remodeling complex with broad regulatory functions across the genome. *CEP290*, a centrosome protein, with essential roles in centrosome and cilia development in many cell types has the second most predicted digenic interactions with 186. The genes with the most predicted digenic pairs were also enriched for several GO metabolic processes. Many of the other genes with large numbers of predicted digenic partners have roles in translation, RNA metabolism and cellular transport (**Figure S17-19**, **Supplementary Tables 5-7**). The top 100 gene pairs with the highest average scores were enriched for electron carriers and transcription co-activators.

We found that 8,649 (26%) of predicted digenic gene pairs had at least one recessive phenotype associated with one of them in OMIM (Amberger et al., 2008, 2015; McKusick, 2007). In more than a quarter of these cases (2241; 25.91%), at least one phenotype was in common or with high semantic similarity (Yujian and Bo, 2007) between the two genes. For most of these gene pairs (2151; 95.98%), there were different OMIM annotation codes, which suggest novel associations.

Many of these predicted novel digenic gene pairs have plausible mechanisms. For example, a digenic pair comprising *STIM1* and *ORAI1* had the 17^th^ highest score over all human gene pairs. It has been previously reported that *STIM1* and *ORAI1* function together to form Ca^2+^ release-activated Ca^2+^ (CRAC) channels, which are responsible for Ca^2+^ influx called store-operated Ca^2+^ entry (SOCE) (Soboloff et al., 2006). The proper functioning of these channels is necessary for maintaining the normal physiology of several cell types, including T cell receptors and human lymphocytes (Lewis, 2001; Lioudyno et al., 2008; Partisetis et al., 1994). Missense variants in *STIM1* and *ORAI1*, individually, cause diseases with a great degree of phenotypic homogeneity (Lacruz and Feske, 2015). Loss of function variants in *STIM1* and *ORAI1* have also been known to cause immunodeficiency, (Feske et al., 2006; McCarl et al., 2009; Picard et al., 2009; Zhang et al., 2015) under autosomal recessive conditions, as reported by OMIM. Therefore, it is possible that single loss of variants in both genes occurring simultaneously could lead to the autosomal recessive immunodeficiency.

## DISCUSSION

In this paper we describe DiGePred, a high-throughput machine-learning approach to identify gene pairs with the potential to cause digenic disease. We demonstrate the accuracy and robustness of our approach in several realistic scenarios. We were motivated to create DiGePred by the challenge of identifying causal variants in patients with rare disease that cannot be explained by a single variant. It is currently unfeasible to experimentally evaluate all potentially causal pairs of variants in a patient of interest. Thus, to facilitate the rapid identification of candidate digenic gene pairs in patients, we provide DiGePred predictions for all pairs of human genes at several confidence thresholds (**Supplementary Tables 4, 8**).

The DiGePred classifier trained using the “unaffected” negative set is best suited to the purpose of identifying digenic pairs in patients with rare disease, because it reflects the baseline distribution of gene pairs with variants in unaffected individuals related to patients using clinical sequencing pipelines. It was also our best performing classifier. However, we demonstrated that our approach preforms well at distinguishing digenic pairs from several different sets of candidate non-digenic gene pairs and that the features used by these classifiers are similar unless the prediction problem is explicitly engineered to make them different (**Figure S6**).

Nonetheless, there is still much to learn about the mechanisms underlying digenic diseases. The features prioritized by our models support previous work (Gazzo et al., 2017, 2016) in that phenotypic similarity, number of phenotypes, and involvement in the same molecular pathways are the most important predictors, and they suggest that these may be more specific predictors of digenic gene pairs than similar co-expression profiles or close interaction network distance. Our results using negatives that match the network and functional features between positives and negatives sets indicate that digenic gene pairs also have differences in their evolutionary attributes. As more digenic disease pairs are identified, we anticipate that even better predictive models will be developed and that these models will yield insight into the genes, pathways, evolutionary histories, and phenotypes associated with digenic pairs.

Our approach intentionally separates the prediction of variants’ effects on gene function from the identification of gene pairs that could cause disease when their functions are disrupted simultaneously. The focus on gene pairs is reflected in our use of gene level and gene-pair level systems biology, biological network, and evolutionary features that represent genes as a whole. The question of whether a variant affects gene function has been studied extensively. There are many methods for interpreting variants of unknown significance,(Adzhubei et al., 2010; Ashkenazy et al.; Celniker et al.; Glaser et al., 2003; Kircher et al., 2014a; Kumar et al., 2009; Rentzsch et al., 2019), but there is low concordance between them (Castellana and Mazza; Dong et al., 2014). The decoupling of these tasks enables users to apply the approaches they believe to be most appropriate for identifying gene pairs of interest before screening for digenic disease potential. For example, in our collaboration with the UDN, this includes application of computational variant effect predictors, study of inheritance patterns, and clinical expertise. Our classifiers perform similarly well whether trained against gene pairs that have predicted disruptive variants or on all variant pairs from individuals (**Figures S11-16**). In the future, it may be beneficial to incorporate variant-level and gene-level information into a single algorithm, in particular in cases where there is structural information about the proteins of interest. Indeed, we have had success incorporating 3D modeling of variants and their interactions with the UDN. However, as we describe in the next paragraph, improper incorporation of variant information has potential to cause high false positive rates.

We compared DiGePred to the recently published ORVAL/VarCOPP digenic disease prediction server. This method was also developed using DIDA as positive training data and is only available as a web server, so it was not possible to evaluate its performance in our training, validation, testing framework. Thus, we applied it to variant gene pairs from the 38 held-out unaffected relatives of UDN patients. This reflects the intended clinical application of the tool. At its strictest (99%) prediction threshold, we found an average of 855 predicted digenic disease pairs per individual without disease. This is an unacceptably high false positive rate for clinical use. In contrast, DiGePred predicts one or fewer digenic pairs for 63% of these individuals and an average of two digenic pairs per individual overall. Our analysis of the ORVAL method suggests that if one of the genes in a pair carries a variant that is predicted to be pathogenic by ORVAL’s variant effect prediction component, then the gene pair is very likely to be predicted to be digenic. Thus, it does not capture a signal specific to digenic disease.

Going forward, there is still much work needed to fully understand and accurately identify novel cases of digenic disease. Most importantly, more characterized examples of digenic diseases and their causal molecular mechanisms are needed. Our analyses are based on the examples available in DIDA, but there are likely hundreds or even thousands of undiscovered cases. We anticipate that our algorithms will further improve with more data. We also believe that there is the potential to integrate information from large-scale screens of genetic and synthetic lethal interactions in human cell lines and model organisms (Gong et al., 2018; Guo et al., 2015; Li et al., 2014; Nijman, 2011; O’Neil et al., 2017; Srivas et al., 2016).

In summary, we have developed DiGePred, a method for identifying gene pairs with digenic disease potential, and generated predictions for all pairs of human genes. Our use of this tool with the UDN illustrates its potential to provide insight in real-world settings, and we anticipate that it will have broad utility in clinical genome interpretation.

## METHODS

### Digenic gene pairs

We obtained known digenic disease gene pairs from the DIgenic Diseases Database (DIDA) (Gazzo et al., 2016). There were 140 unique gene pairs in DIDA. These pairs served as the “positive” training data for the machine learning classifier and were termed the *digenic* set of gene pairs. DIDA provides information about the genes mutated together in cases of digenic disease, the variants in the genes, the number of variants on both alleles, as well as information concerning the connectivity of the genes forming a gene pair such as distance on PPI network, whether expressed in same tissue, whether members of the same biochemical pathway, and whether annotated to have the same function. The additional list of digenic pairs discussed in a follow up paper by the group that produced DIDA (Gazzo et al., 2017) were not used for training.

### Non-digenic gene pairs

We generated several sets of putative non-digenic gene pairs that served as the “negative” data in training different classifiers. The *unaffected* non-digenic set was created from genes with variants in the sequenced exomes or genomes of relatives of UDN patients deemed unaffected by the UDN. Thus, we consider any combination of genes observed to be mutated simultaneously in any one “unaffected” individual to be non-digenic. Combining gene pairs from 55 individuals, the unaffected set contains 1.8 million gene pairs. The *random* non-digenic set was created by selecting random pairs from the list of all human genes. The *permuted* non-digenic set was created by generating all possible pairs of two genes from the DIDA genes excluding actual DIDA pairs; this resulted in 13,390 permuted gene pairs. We created the *matched* non-digenic gene pair set from the random gene pairs by selecting gene pairs such that the distribution of the six NFFs match those of the digenic set. The digenic gene pairs were binned by dividing the distribution of features into equal sized intervals, such that every feature value data interval had an equal number of gene pairs. We selected random gene pairs for the matched set such that the distributions of feature values for all the selected pairs recapitulated the overall distribution for all features of the digenic set, simultaneously.

### Six Network and Functional Features

#### Pathway similarity

The pathway annotations for the genes were derived from KEGG (Kanehisa et al., 2017) and Reactome (Fabregat et al., 2018). The Jaccard similarity metric (Jaccard, 1912) was used to calculate the proportion of pathway overlap between the two genes. The Jaccard similarity is measured by the ratio between the intersection of two sets and the union of two sets. In this case, the pathway similarity was calculated by taking the ratio of pathways annotations in common with both genes and pathway annotations associated with either. If both genes did not have pathway annotation, the similarity value was 0.

#### Phenotype similarity

The phenotype annotations from the Human Phenotype Ontology (HPO) (Köhler et al., 2017) for the genes were used as features. The phenotypic overlap between the two genes was calculated similarly, as above, using the Jaccard similarity metric. The value for missing phenotype annotations was 0.

#### Co-expression

The co-expession data was derived from the COXPRESdb web server version 7.3 (Okamura et al., 2015). The data is in the form of a mutual co-expression rank, which indicated how likely it was for a pair of genes to be co-expressed in the same tissue and the same level compared to other gene pairs. A lower rank indicated high co-expression. The inverse of the rank was used as the feature and if either gene was not found in the co-expression database, the value used was 0.

The network data was downloaded from the UCSC gene and pathway interaction browser (Poon et al., 2014), which in turn was derived from other sources of data, such as protein-protein interaction (PPI) databases (Ruepp et al., 2007, 2010; Szklarczyk et al., 2017; Turner et al., 2010), functional annotation databases (Nédélec et al., 2016) and others.

#### PPI distance

The PPI network was based on experimental data regarding protein interactions. The inverse of the shortest path between a pair of genes on this network was used as the PPI distance feature.

#### Pathway distance

The pathways interaction network was based on interactions between the various curated biochemical pathways. The inverse of the shortest path between a pair of genes on this network was used as the pathway distance feature.

#### Literature distance

The literature mined interaction network was made up of interactions derived from reported interactions or predicted associations in published biomedical literature. The inverse of the shortest path between a pair of genes on this network was used as the literature distance feature.

### Five Evolutionary Features

#### Evolutionary age

We obtained the evolutionary ages of the proteins coded by the genes using ProteinHistorian (Capra et al., 2012). This conveyed the idea of how long ago did they first evolve and in which organisms. Older genes are usually more conserved and dysfunction of these genes could have a considerable impact on normal physiology. The scores for both genes were used.

#### Gene essentiality

The gene essentiality scores provide a rank of how important and vital a gene is for normal physiology, viability and survival. They were derived from the OGEE webserver (Chen et al., 2012, 2017). The essentiality scores are based on knockout (KO) experiments in model organisms and cell based assays. The scores for both genes were used.

#### Loss of function intolerance (pLI)

We added the loss of function intolerance (pLI) scores (Fadista et al., 2016), obtained from the EXAC consortium. These scores were based on the difference between actual mutation incidence and expected mutation frequency. A depletion of mutation incidence, compared to expected frequency, could mean the inability of the organism to survive if the gene was mutated. The scores for both genes were used.

#### Selection pressure (dN/dS)

We used measures of selection pressure in the form of dN/dS scores for the genes. These were derived from the EVOLA web server (Matsuya et al., 2007). dN/dS ratios give a measure of the ratio between the non-synonymous mutations and synonymous mutations during evolution. This ratio tells us whether the gene has been evolving under strong positive, negative or neutral selection. The scores for both genes were used.

#### Haploinsufficiency

We used the Haploinsufficiency scores (Huang et al., 2010) which were in the form of predictions of which genes were haploinsufficient, based on observed mutations. The scores for both genes were used.

### Gene-focused network and functional features

#### Number of pathways

The features used for the classifier were the number of pathways associated with gene A and the number of pathways associated with gene B

#### Number of phenotypes

Similar to the pathways, the features used for the classifier were number of phenotypes associated with gene A and with gene B, individually.

#### Network neighbors

The features used were number of genes in the network, directly connected to gene A and to gene B, individually. Additionally, the number of shared network neighbors was also used which was defined as the number of genes directly connected to both gene A and B. These metrics were defined for all three types of interaction networks.

#### Number co-expressed

The features used in the classifier were number of genes highly co-expressed with gene A and the number of genes highly co-expressed with gene B, individually. Additionally, the number of genes highly co-expressed with both gene A and gene B, was used. Highly co-expressed genes were defined using a mutual co-expression rank of 500 (out of possible 20,000).

### Performance Quantification

Receiver Operating Characteristic (ROC) and Precision-Recall (PR) curves were computed to evaluate the performance of the classifiers. The ROC curve plots the False Positive Rate (FPR) on the x-axis and the True Positive Rate (TPR) on the y-axis. The area under each curve (AUC) was used to summarize performance.

### Training and Testing the DiGePred Random Forest Models

We trained several random forest (RF) classifiers to distinguish digenic and non-digenic gene pairs. We selected RFs because they can integrate diverse features, perform well on unbalanced positive and negative sets, and provide interpretable models. The sci-kit learn (sklearn) python module was used for all training, evaluation, and prediction (Pedregosa et al., 2011). Hyper-parameters were selected by nested cross validation on 80% of the labeled gene pairs. A stratified shuffle split was used for 10-fold cross validation. This method involved splitting the data into 10 equal parts, with each part of the data containing approximately the same ratio of positives and negatives as the other parts. The optimum number of trees was found to be 500 and the maximum depth was found to be 15. Based on these analyses, we selected the classifier trained with the unaffected negative pairs and all features as the best model, and we refer to this as the DiGePred classifier.

The remaining 20% of the combined labeled data was held out for final performance validation of this best model from the cross-validation. These pairs had not been previously evaluated by the classifier. In addition to the held-out positive digenic pairs, we generated 100 sets of held-out non-digenic pairs for evaluation. This enabled us to evaluate the best classifier 100 fold, with the same positive digenic pairs used in every iteration, but a unique non-overlapping set of held-out non-digenic pairs in every iteration.

### Evaluation using additional digenic pairs not in DIDA

The classifier was further evaluated using an external set, made up of gene pairs considered to be digenic that were reported after DIDA was compiled. The external evaluation set was used in the previously published variant combination pathogenicity predictor (VarCOPP/ ORVAL) (Papadimitriou et al., 2019; Renaux et al., 2019). This set had three unique gene pairs, which did not overlap with DIDA pairs. These gene pairs (AHI1, CEP290), (CEP290, CRB1) and (CEP290, RPE65) was labeled Papadimitrou et al., 19 validation set. We included recently discovered novel digenic inheritance of profound non-syndromic hearing impairment caused by (PCDH15, USH1G) (Schrauwen et al., 2018). In addition, three recently reported cases of digenic inheritance in immune disorders were used. Ameratunga et al., 17 identified epistatic interactions between TACI and TCF3 (or TNFRSF13B) resulting in severe primary immunodeficiency disorder and systemic lupus erythematosus (Ameratunga et al., 2017). Hoyos-Bachiloglu et al., 17 discussed how human immunodeficiency was caused by mutations in IFNAR1 and IFNGR2 (Hoyos-Bachiloglu et al., 2017). More recent digenic findings such as (LAMA4, MYH7) linked to infantile dilated cardiomyopathy (Abdallah et al., 2019) from Abdallah et al., 19; (KCNE2, KCNH2) linked to long QT syndrome types 2 and 6 (Heida et al., 2019) from Heida et al., 2019; (CLCNKB, SLC12A3) linked to Gitelman syndrome (Kong et al., 2019) from Kong et al., 2019; (CACNA1C, SCN5A) linked to Long QT phenotype (Nieto-Marín et al., 2019) from Nieto-Marín et al., 2019; (FGFR1, KLB) linked to insulin resistance (Stone et al., 2019) and diabetes from Stone et al., 2019; (CLCNKA, CLCNKB) linked to Bartter syndrome with sensorineural deafness (Nozu et al., 2008) from Nozu et al., 2008; and (CLCN7, TCIRG1) linked to osteoporosis (Yang et al., 2018) from Yang et al., 2018 were used to assess the classifier as well.

We also included gene pairs not characterized as digenic, but displaying functional synergy associated with disease or adverse phenotypes. We derived the gene pair from the previously reported UDN study that found mutations in TRPS1 and FBN1 to be responsible for the patient phenotype and it was labeled *Zastrow et al., 17 (UDN))* (Zastrow et al., 2017).

### Feature Importance

To identify the most indicative features we used the inbuilt classifier feature importance function in sklearn, which uses the Gini impurity approach to quantify the relative feature importance for all features.

### Prediction Score Thresholds

We determined a digenic score threshold for the DiGePred classifier for classifying gene pairs digenic based on the F_0.5_ metric. This is a modification of the F_1_ statistic, designed to attenuate the effect of false negatives. It is calculated as *F_ß_ = (1 + ß2) × TP / (1 + ß2 × TP + ß2 × FP + FP)*, where ß=0.5, TP=true positives, FP=false positives. The score that yielded the highest F_0.5_ value was 0.534.

### Estimating the False Positive Rate at various score thresholds

We evaluated the DiGePred classifier with an external set of non-digenic gene pairs as well. These gene pairs were obtained from 38 unaffected relatives of UDN patients. The genes were preliminarily selected if the variant in the gene had an ExAC (Lek et al., 2016; 2016) minor allele frequency of < 1%. A gene was further selected if it received a pathogenicity score of ‘D’ *(“probably damaging”*) from Polyphen2 (Kircher et al., 2014) Only genes passing this Polyphen2 filter were selected to limit the predictions to pairs of genes with variants that likely affected molecular function.

Additionally, genes with rare variants were selected based on a consensus pathogenicity approach if at least two out of Polyphen2, SIFT (Sim et al., 2012; Vaser et al., 2015), CADD (Kircher et al., 2014; Rentzsch et al., 2019), Omicia (Coonrod et al., 2013) and PhyloP (Pollard et al., 2010) agreed that the variant(s) in the gene was pathogenic. A Polyphen 2 selection criteria was similar to before. A variant was deemed pathogenic by SIFT if the score was <=0.05. a CADD score >= 30 was considered pathogenic, while a PhyloP score of <= –10 or an Omicia score >= 0.93 for a variant deemed it pathogenic. All possible gene pairs were used as the consensus pathogenic gene pairs for an individual.

### Comparison with ORVAL

We submitted the list of gene pairs for all the unaffected individuals to the ORVAL(Papadimitriou et al., 2019; Renaux et al., 2019) server. We compared the number of pairs predicted to be digenic by ORVAL, according to its highest confidence threshold, to the number predicted by our method to be digenic at the F_0.5_ threshold. We obtained the list of genes for each unaffected individual as mentioned in the previous section. We evaluated the statistical significance of the number of digenic pairs predicted as false positives per individual between DiGePred and ORVAL using a MWU test.

Furthermore, 20% of all genes with rare variants in the individual were chosen at random. All possible gene pairs were generated to constitute the random set of gene pairs for each individual. We calculated the number of digenic pairs predicted per individual at different score thresholds. This was done to compare the number of false positives between ORVAL and DiGePred fairly. As ORVAL includes variant effects as a feature, selecting for genes with variants that were predicted pathogenic by Polyphen2 or by a consensus of several predictors of variant effect could bias against ORVAL, though it reflects common clinical practice. Therefore, we also compared DiGePred and ORVAL on pairs of genes selected at random.

### Gene ontology (GO) enrichment

The GO enrichment was computed using a web resource WebGestalt (WEB-based GEne SeT AnaLysis Toolkit) (Liao et al., 2019). A list of genes was prepared for each selected set of predicted digenic pairs based on highest score, highest average score or most predicted pairs. This list of genes was ranked based on the selection criteria and the GO enrichment for biological process, cellular component and molecular function categories.

## ACKNOWLEDGEMENTS

This work was supported by an award from the National Institutes of Health (NIH) Common Fund, through the Office of Strategic Coordination and the Office of the NIH Director, to the clinical sites (U01HG007674, to Vanderbilt University Medical Center). It was also supported by NIH award (R35GM127087 to JAC). This work was conducted in part using the resources of the Advanced Computing Center for Research and Education at Vanderbilt University, Nashville, TN. We thank members of the Capra, Meiler, and UDN Labs for helpful comments. Finally, we thank all patients and their families.

