## Supplementary Tables for "Identifying digenic disease genes using machine learning in the undiagnosed diseases network"

Table available upon request

**SUPPLEMENTARY TABLE 1: HELD-OUT DIGENIC GENE PAIRS**

DiGePred predictions on held-out digenic gene pairs from DIDA (n=28). These pairs were not used for training and used to test the trained classifier.

| # | Gene A | Gene B | Paper | Phenotypes | DiGePred score |
|---|---|---|---|---|---|
| **1** | AHI1 | CEP290 | Coppieters et al., 10, Cheng et al., 12<br>Papadimitriou et al., 19 (validation set) | Leber Congenital Amaurosis, Joubert syndrome | 0.756 |
| **2** | CEP290 | RPE65 | Coppieters et al., 10 | Leber Congenital Amaurosis | 0.747 |
| **3** | CEP290 | CRB1 | Coppieters et al., 10 | Leber Congenital Amaurosis | 0.676 |
| **4** | TCF3 | TNFRSF13B | Ameratunga et al., 17 | Primary immunodeficiency disorder and systemic lupus erythematosus | 0.637 |
| **5** | IFNAR1 | IFNGR2 | Hoyos-Bachiloglu et al., 17 | Primary immunodeficiency | 0.131 |
| **6** | PCDH15 | USH1G | Schrauwen et al., 17 | Profound non-syndromic hearing impairment | 0.769 |
| **7** | LAMA4 | MYH7 | Abdallah et al., 19 | Infantile Dilated Cardiomyopathy | 0.347 |
| **8** | KCNE2 | KCNH2 | Heida et al., 19 | Long QT Syndrome Type 2 and Type 6 | 0.848 |
| **9** | CLCNKB | SLC12A3 | Kong et al., 19 | Gitelman syndrome | 0.636 |
| **10** | CACNA1C | SCN5A | Nieto-Marín et al., 19 | Long QT phenotype | 0.516 |
| **11** | FGFR1 | KLB | Stone et al., 19 | Endocrine Specific FGF-21 Signaling Defects and Extreme Insulin Resistance | 0.45 |
| **12** | CLCN7 | TCIRG1 | Yang et al., 18 | Osteopetrosis | 0.723 |
| **13** | CLCNKA | CLCNKB | Nozu et al., 2008 | Bartter syndrome, sensorineural deafness | 0.744 |
| ***14*** | *FBN1* | *TRPS1* | *Zastrow et al., 17 (UDN)* | *Marfan syndrome and TRPS1* | *0.195* |

**SUPPLEMENTARY TABLE 2: NOVEL DIGENIC PAIRS NOT IN DIGENIC DATABASE (DIDA) FROM RECENT LITERATURE**

Genes, publication, and patient phenotypes for 13 recently identified digenic disease pairs. Red indicates a DiGePred score beneath the F0.5 threshold. The 14$^{th}$ pair (italics) is not strictly digenic, but underlies disease in a UDN patient.

Table available upon request

**SUPPLEMENTARY TABLE 3: NOVEL DIGENIC GENE PAIRS FROM LITERATURE**

DiGePred predictions on novel digenic gene pairs from recent literature (n=13). These pairs were not included in DIDA.

Table available upon request

**SUPPLEMENTARY TABLE 4: PREDICTED DIGENIC PAIRS FROM ALL POSSIBLE HUMAN GENE PAIRS**

Gene pairs predicted to be digenic by DiGePred at most confident threshold. (n=33,272)

| GO term | Description | Size | Enrichment score | Normalized enrichment score | P-Value | FDR |
|---|---|---|---|---|---|---|
| GO:0003735 | structural constituent of ribosome | 21 | 0.73 | 1.95 | ~0 | 0.0009 |
| GO:0006413 | translational initiation | 20 | 0.71 | 1.92 | ~0 | 0.0017 |
| GO:0070972 | protein localization to endoplasmic reticulum | 20 | 0.71 | 1.92 | ~0 | 0.0017 |
| GO:0006401 | RNA catabolic process | 21 | 0.70 | 1.89 | ~0 | 0.0017 |
| GO:0090150 | establishment of protein localization to membrane | 22 | 0.69 | 1.88 | 0.001 | 0.0018 |
| GO:0006605 | protein targeting | 22 | 0.70 | 1.9 | ~0 | 0.0022 |

**SUPPLEMENTARY TABLE 5: GENE ONTOLOGY ENRICHMENT FOR TOP 100 GENES WITH MOST PREDICTED DIGENIC PAIRS**

Gene ontology (GO) enrichment using WebGestalt (WEB-based GEne SeT AnaLysis Toolkit).

| GO term | Description | Size | Enrichment score | Normalized enrichment score | P-Value | FDR |
|---|---|---|---|---|---|---|
| GO:0009055 | electron transfer activity | 6 | -0.883 | -2.299 | 0 | 0 |
| GO:0006091 | generation of precursor metabolites and energy | 8 | -0.902 | -2.73 | 0 | 0 |
| GO:0030139 | endocytic vesicle | 5 | -0.758 | -1.83 | 0 | 0.0095 |

**SUPPLEMENTARY TABLE 6: GENE ONTOLOGY ENRICHMENT FOR TOP 100 GENES WITH HIGHEST AVERAGE PREDICTED VALUE**

Gene ontology (GO) enrichment using WebGestalt (WEB-based GEne SeT AnaLysis Toolkit).

| GO term | Description | Size | Enrichment score | Normalized enrichment score | P-Value | FDR |
|---|---|---|---|---|---|---|
| GO:0097730 | non-motile cilium | 9 | 0.695 | 1.72 | 0.0055 | 0.02 |
| GO:0045111 | intermediate filament cytoskeleton | 6 | 0.779 | 1.72 | 0.0023 | 0.037 |
| GO:0043473 | pigmentation | 6 | -0.953 | -2.52 | 0 | 0 |
| GO:0045444 | fat cell differentiation | 7 | -0.918 | -2.55 | 0 | 0 |

**SUPPLEMENTARY TABLE 7: GENE ONTOLOGY ENRICHMENT FOR GENES IN THE TOP 100 GENE PAIRS WITH HIGHEST PREDICTED VALUE**

Gene ontology (GO) enrichment using WebGestalt (WEB-based GEne SeT AnaLysis Toolkit).

Table available upon request

**SUPPLEMENTARY TABLE 8: DIGENIC PREDICTIONS ON ALL HUMAN GENE PAIRS**

DiGePred predictions on all possible human gene pairs (n=155.32 mil.)
