## Supplementary Figures for "Identifying digenic disease genes using machine learning in the undiagnosed diseases network"

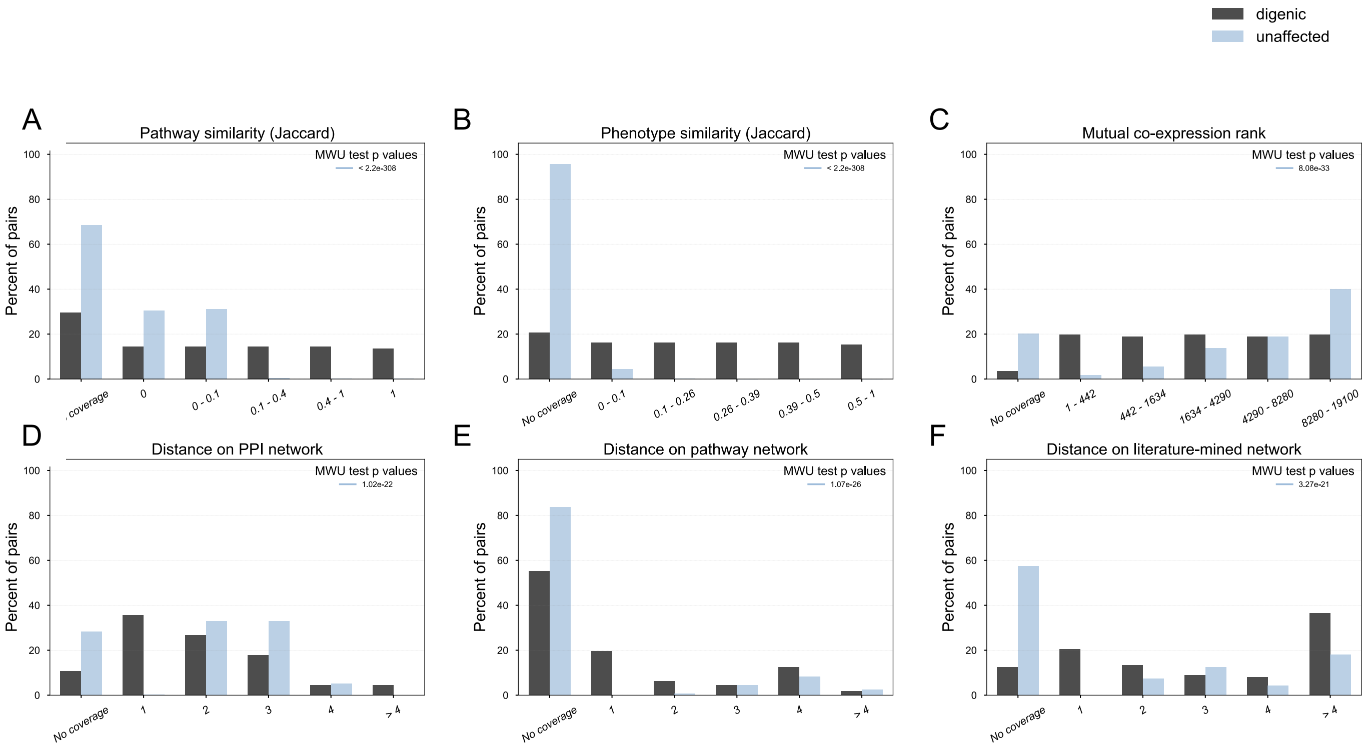


**FIGURE S1: DISTRIBUTION OF NETWORK FEATURES IS DIFFERENT FOR DIGENIC VS. UNAFFECTED GENE PAIRS**

Feature distributions of network and functional features (NFFs) for digenic gene pairs (black) and unaffected gene pairs (blue). Feature value bins are shown along the X-axis and the proportion of the gene pair set on the Y-axis. Distributions were compared using the Mann-Whitney U (MWU) test. A) Jaccard similarity of KEGG and Reactome pathways associated with both genes; B) Jaccard similarity of HPO phenotypes associated with both genes; C) Mutual co-expression rank of gene pair compared to all other gene pairs; D-F) Distance on experimental protein-protein interaction (PPI), biochemical pathways, and literature-mined interaction networks, obtained from UCSC gene and pathway interaction browser database. All six NFFs are significantly different between the gene pair sets (MWU $P < 10^{-20}$).

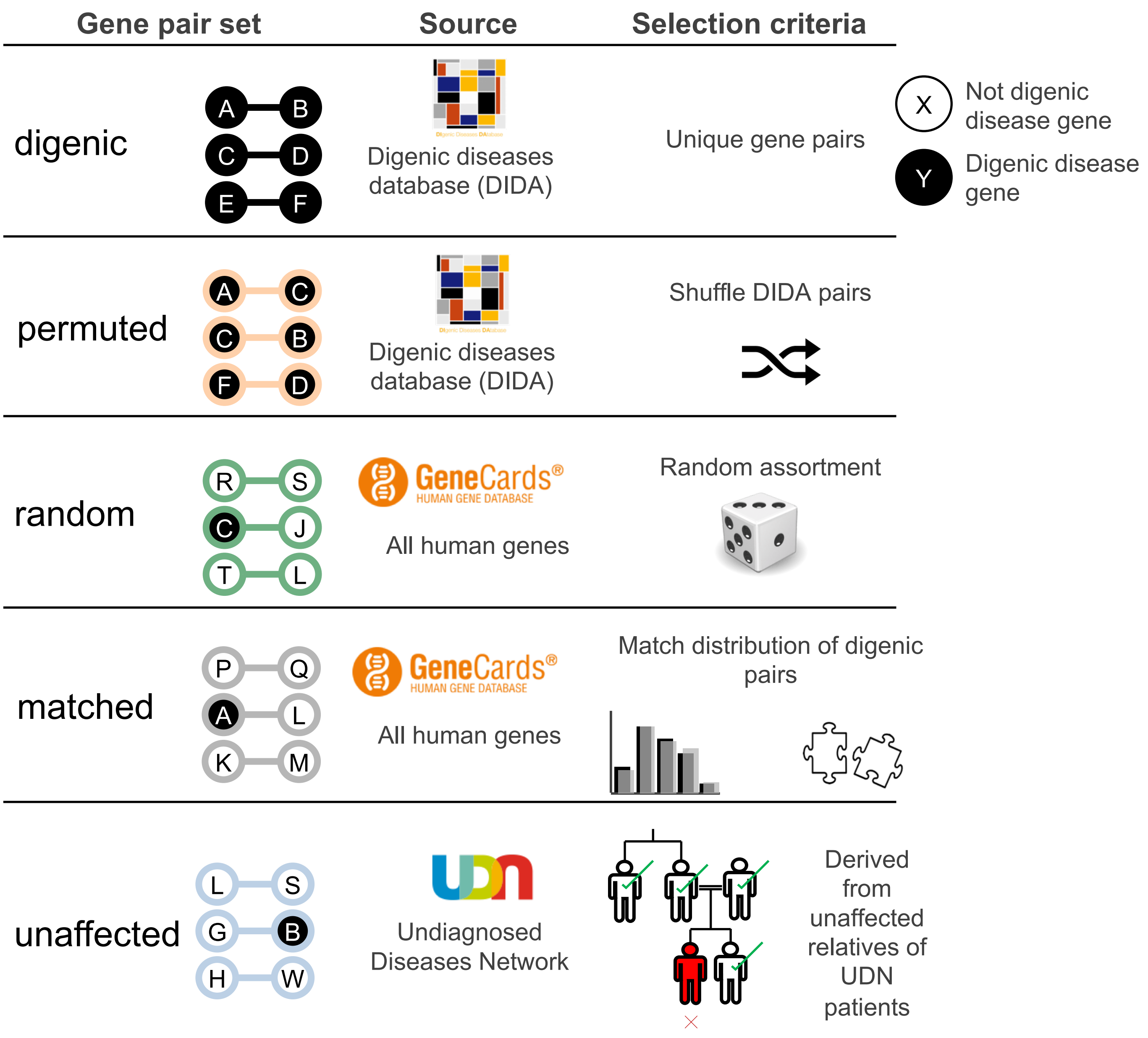


**FIGURE S2: SCHEMATIC OF ALL POSITIVE AND NEGATIVE TRAINING SETS USED FOR CLASSIFICATION OF DIGENIC DISEASE GENES**

Digenic gene pairs were derived from the Digenic Diseases Database (DIDA). Unique digenic gene pair combinations (n=140) were used for training and evaluation. Permuted negative gene pairs were generated by computing all possible permutations of genes in digenic pairs, excluding the known digenic combinations. Random gene pairs were generated by selecting random pairs of all human genes, excluding any known to be digenic. Matched gene pairs were selected from random gene pairs so that the set matched the NFF distribution of digenic pairs (**Figure S1**). Unaffected gene pairs were derived from genes with variants in unaffected members of UDN patient families.

Evolutionary features

A

| Feature | Source | Logic |
|---|---|---|
| loss of function intolerance<br>haploinsufficiency | DECIPHER GRCh37 | 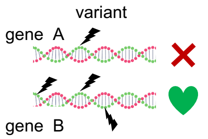  Ability of gene to withstand mutations |
| protein age | ProteinHistorian | 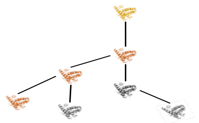 Evolutionary history of protein coded by gene |
| selection pressure | Evola | 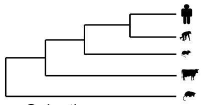 Selection pressures acting on gene during evolution |
| gene essentiality | OGEE | 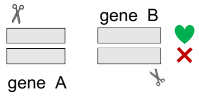  Gene essentiality derived from gene knockout experiments |

Gene focused network and functional features

B

| Feature | Related NFF |
|---|---|
| Number of pathways gene A<br>Number of pathways gene B | pathway similarity |
| Number of phenotypes gene A<br>Number of phenotypes gene B | phenotype similarity |
| Number of highly co-expressed with gene A<br>Number of highly co-expressed with gene B<br>Number common highly co-expressed | co-expression rank |
| Number of neighbors gene A<br>Number of neighbors gene B<br>Neighbor similarity | PPI distance<br>pathway distance<br>literature distance |

**FIGURE S3: ADDITIONAL FEATURE SETS USED FOR MACHINE LEARNING CLASSIFICATION OF DIGENIC DISEASES**

**(A)** Individual gene level Evolutionary biology and genomics features: i) *loss of function intolerance*; ii) *haploinsufficiency*, measures of mutational load on gene; ii) *protein age*, measure of evolutionary age of protein coded by gene; iv) *dN/dS score*, measures the constraints on selection during mammalian evolution of gene; v) *essentiality score*, derived from gene KO experiments, measures how vital gene is to organism survival. **(B)** Individual gene level NFF related features: i) number of pathways associated with both genes; ii) number of phenotypes with both genes; iii) number of genes highly co-expressed with both genes; iv) number of neighbors on biochemical interaction networks, for both genes.

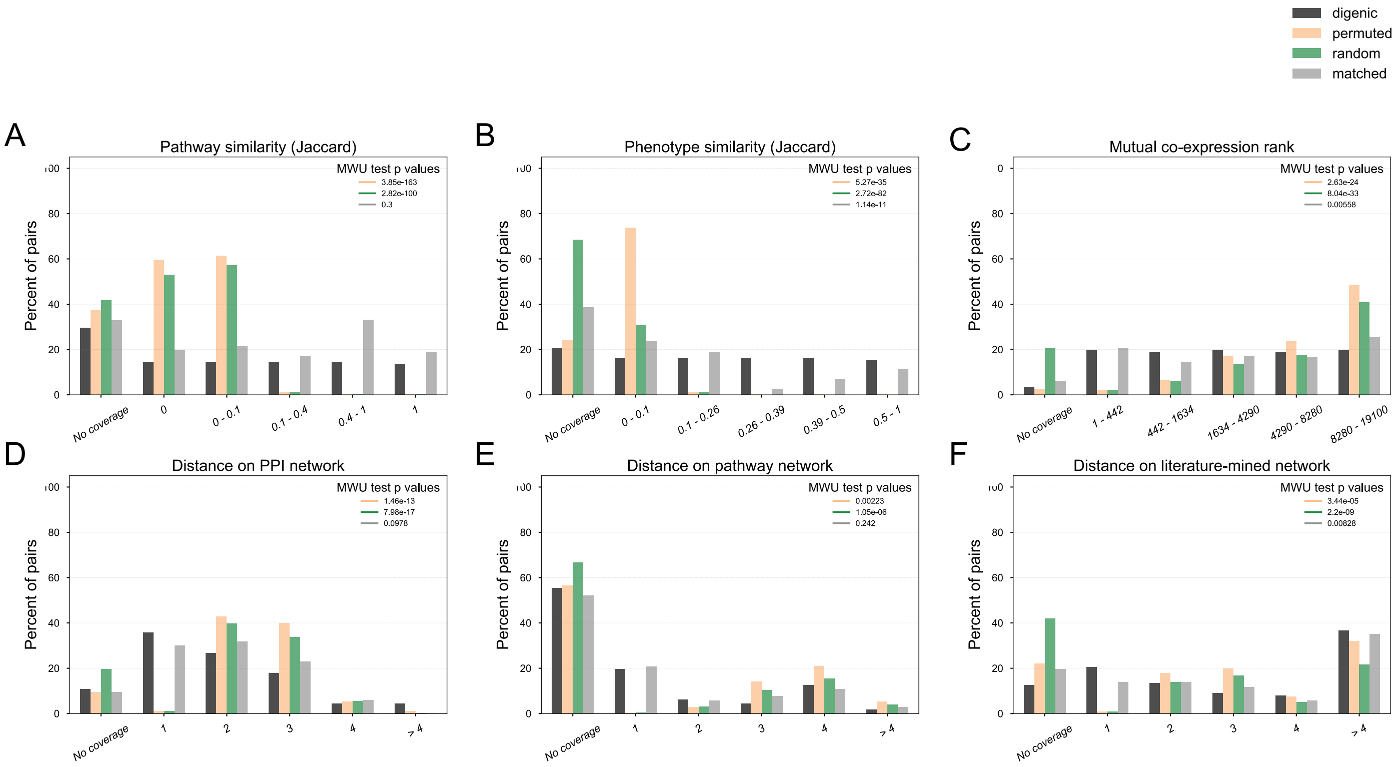


**FIGURE S4: DISTRIBUTION OF NETWORK FEATURES IS DIFFERENT FOR DIGENIC AND NON-DIGENIC PAIRS; SIMILAR FOR MATCHED GENE PAIRS**

Feature distributions of network and functional features (NFFs) for digenic gene pairs (black), permuted (orange), random (green) and matched (grey) gene pairs. The feature value bins shown along X axis and proportion of gene pairs along Y axis. Distributions compared using MWU test, P values shown. **A)** Jaccard similarity of KEGG and Reactome pathways associated with both genes; **B)** Jaccard similarity of HPO phenotypes associated with both genes; **C)** Mutual co-expression rank, comparison of co-expression of gene pair to all other gene pairs across multiple co-expression platforms; **D-F)** Distance on experimental PPI, biochemical pathways and literature-mined interaction networks, obtained from UCSC gene and pathway interaction browser database. MWU P < $10^{-4}$ for permuted and random gene pairs but lower for matched.

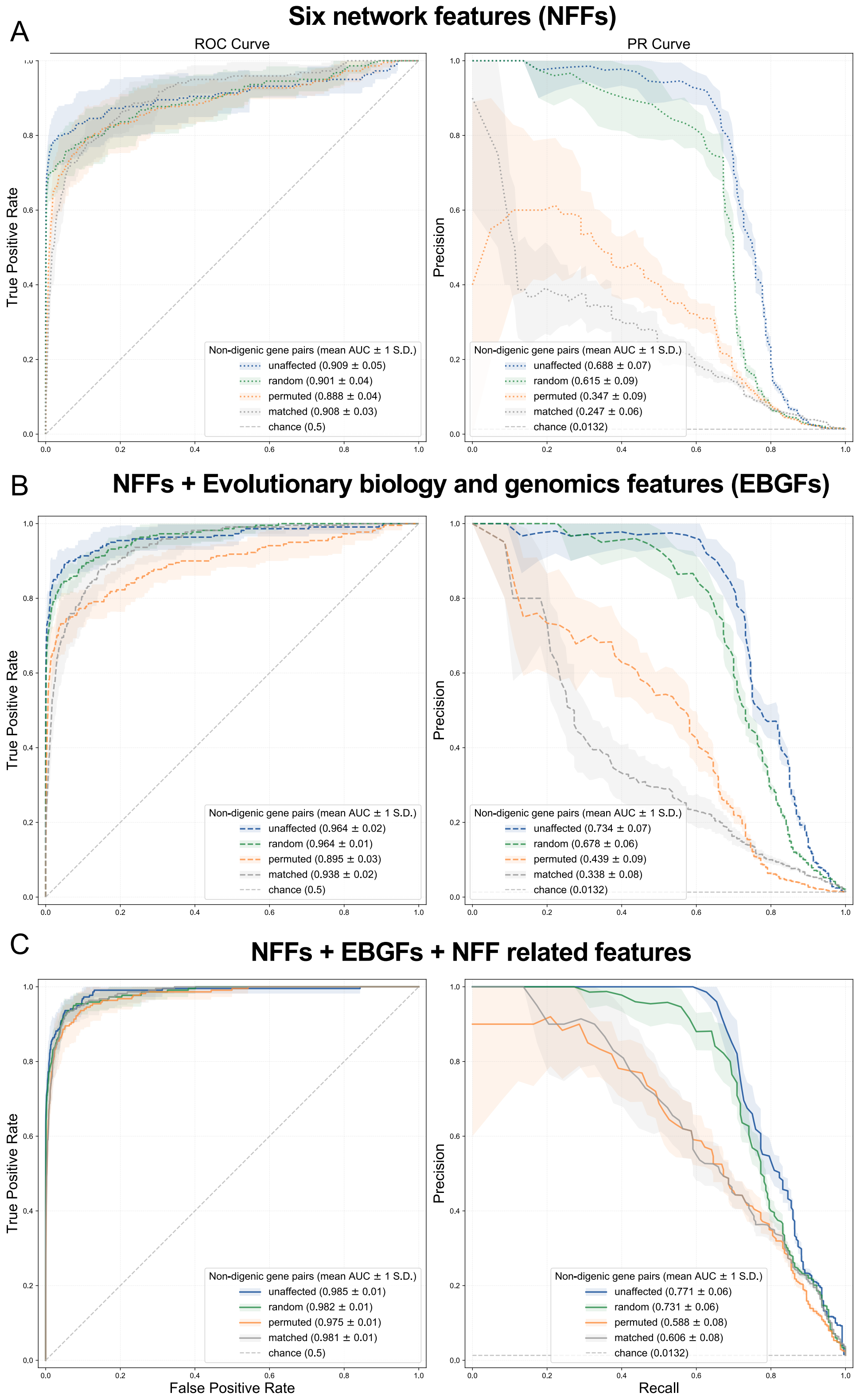


**FIGURE S5: CLASSIFIER ACCURATELY IDENTIFIED DIGENIC PAIRS FROM ALL NON-DIGENIC GENE PAIRS USING VARIOUS FEATURE SETS; ADDITION OF FEATURES IMPROVED PERFORMANCE**

Performance of the classifier on distinguishing between digenic pairs and non-digenic sets of gene pairs: **U**naffected (blue), **R**andom (green), **P**ermuted (orange), **M**atched (grey) training data, measured by area under the Receiver Operating Characteristic (ROC) and Precision-Recall (PR) curves (AUCs), using different feature sets: **A)** six network features (NFFs) (dotted line) AUCs=0.91/0.69U, 0.90/0.62R, 0.88/0.35P, 0.91/0.25M ; **B)** NFFs + Evolutionary biology and genomics features (EBGFs) (dashed line) AUCs=0.97/0.73U, 0.97/0.68R, 0.90/0.44P, 0.94/0.34M; **C)** NFFs + EBGFs + NFF related features (solid line) AUCs=0.99/0.77U, 0.98/0.73R, 0.98/0.59P, 0.95/0.61M.

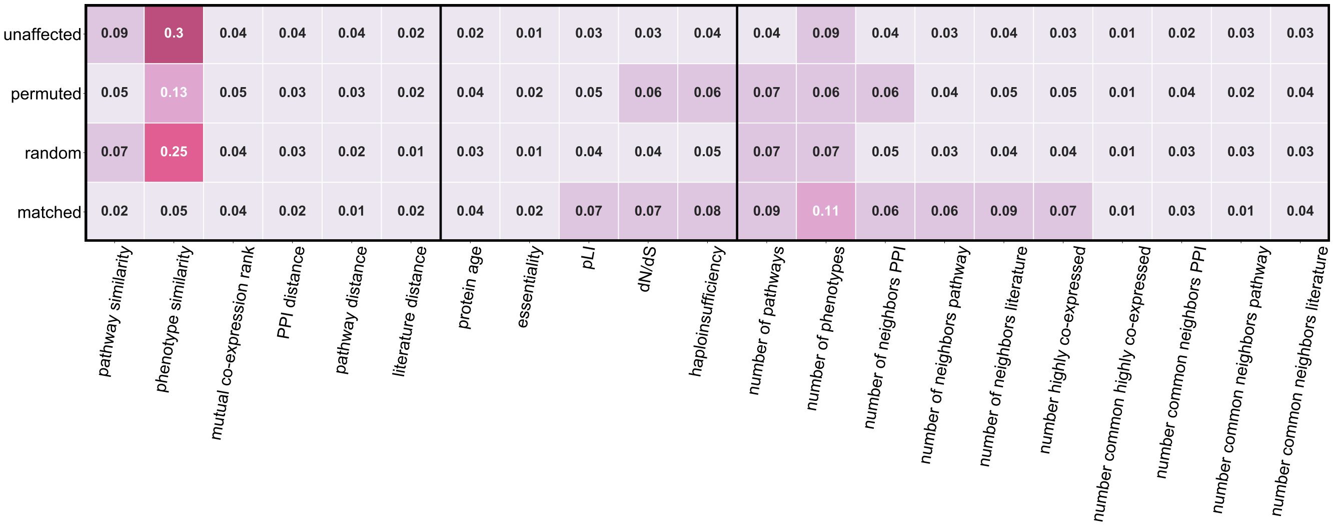


**FIGURE S6: PHENOTYPE FEATURES WERE MOST IMPORTANT FOR CLASSIFIER TO IDENITFY DIGENIC PAIRS; EVOLUTIONARY FEATURES MORE IMPORTANT FOR MATCHED PAIRS**

GINI feature importance values for classifier to identify digenic gene pairs from unaffected, random, matched and permuted non-digenic gene pairs. Phenotype similarity and number of phenotypes had highest importance (> 30%), followed by pathway similarity (~9%). Evolutionary and genomics features more important when distinguishing digenic pairs from matched gene pairs.

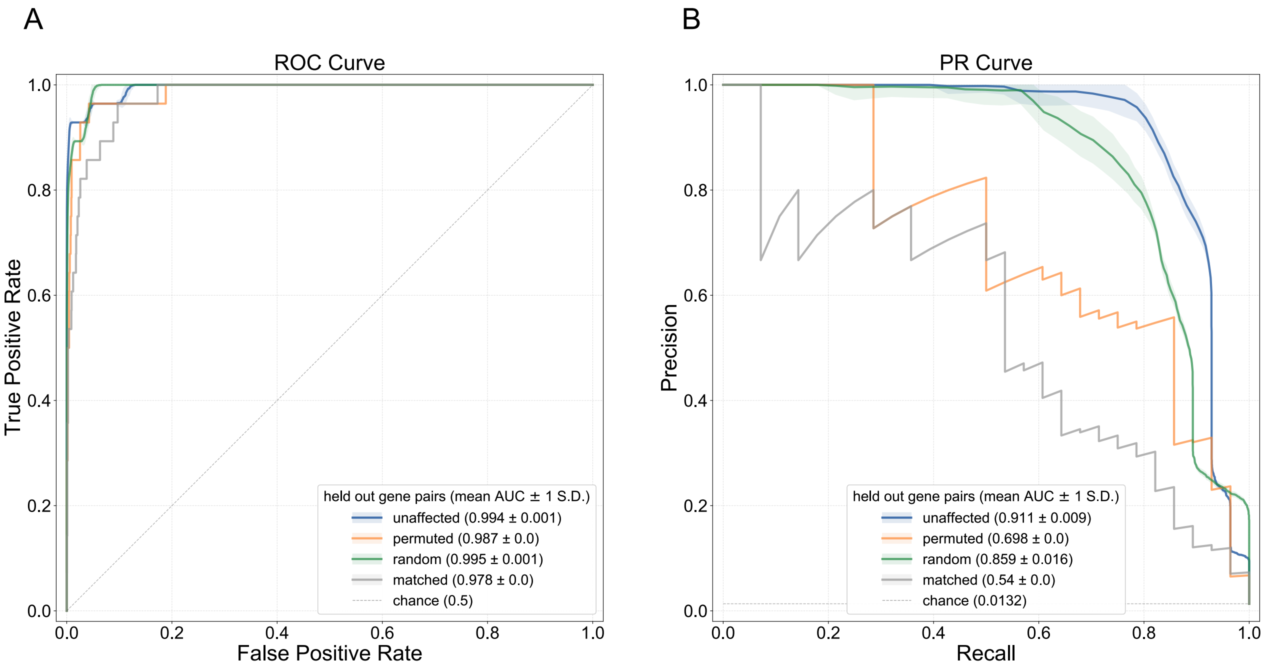


**FIGURE S7: CLASSIFIER TRAINED ON ALL NON-DIGENIC GENE PAIRS ACCURATELY IDENTIFIED DIGENIC PAIRS IN HELD-OUT SET**

Performance of DiGePred (trained on all features using the various negative sets) on the the corresponding held-out test set as evaluated by **(A)** ROC and **(B)** PR curves. We considered four different negative sets: i) *Unaffected*, derived from healthy relatives of UDN patients (blue); ii) *Random*, derived by randomly selecting pairs of genes (green); iii) *Permuted*, derived by generating permutations of known digenic pairs (orange); iv) *Matched*, derived by matching the distribution of network and functional features observed among the digenic pairs (grey). The area under the ROC curves (AUROCs) were > 0.97 in all cases, while the area under the PR curve (AUPRs) were > 0.5 in all cases. These values were much higher than what is expected by chance.

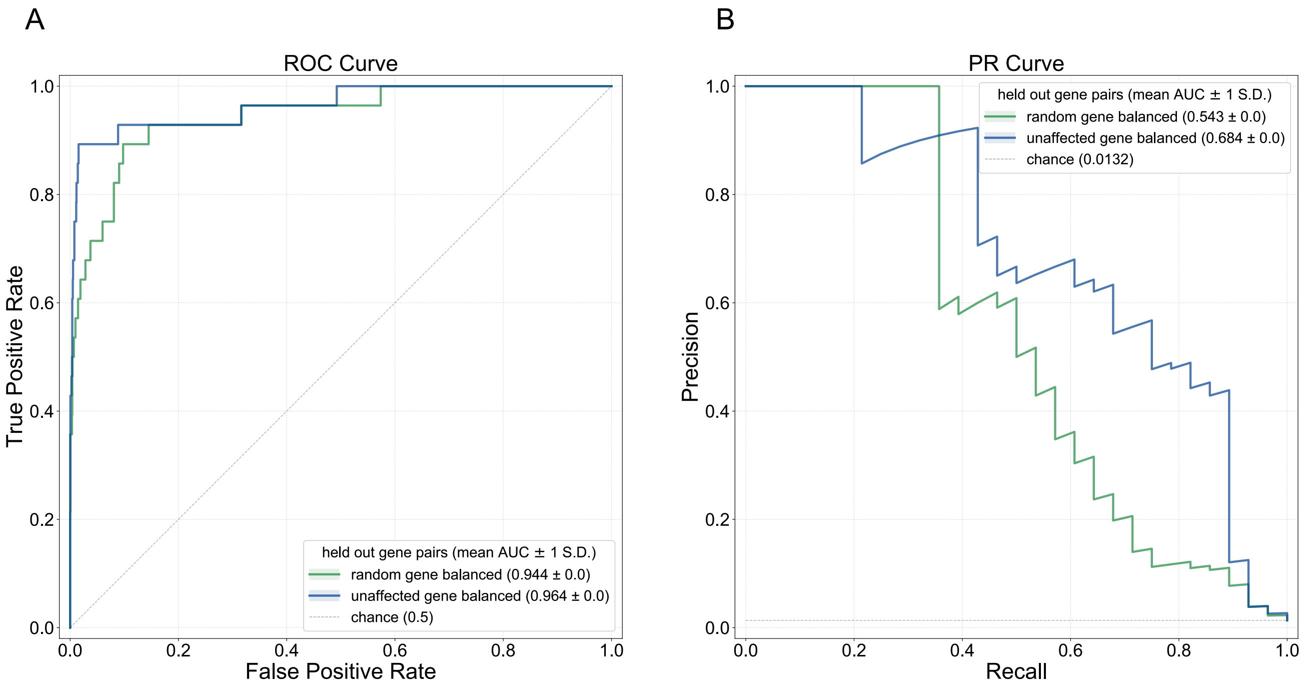


**FIGURE S8: CLASSIFIER ACCURATELY IDENTIFIED DIGENIC PAIRS IN HELD-OUT SET IN THE CASES WEHRE THERE WAS NO GENE OVERLAP BETWEEN TRAINING AND TESTING DATASETS**

Performance of DiGePred (trained on all features using the gene balanced negative sets where there were no overlapping genes between the training and testing datasets) on the corresponding held-out test set as evaluated by **(A)** ROC and **(B)** PR curves. We considered two different gene balanced negative sets: i) *Unaffected gene balanced*, derived from healthy relatives of UDN patients and no genes in common between the training and testing datasets (blue); ii) *Random gene balanced*, derived by randomly selecting pairs of genes no genes in common between the training and testing datasets (green); The area under the ROC curves (AUROCs) were >0.94 in both cases, while the area under the PR curve (AUPRs) were >0.54 in all cases. These values were much higher than what is expected by chance.

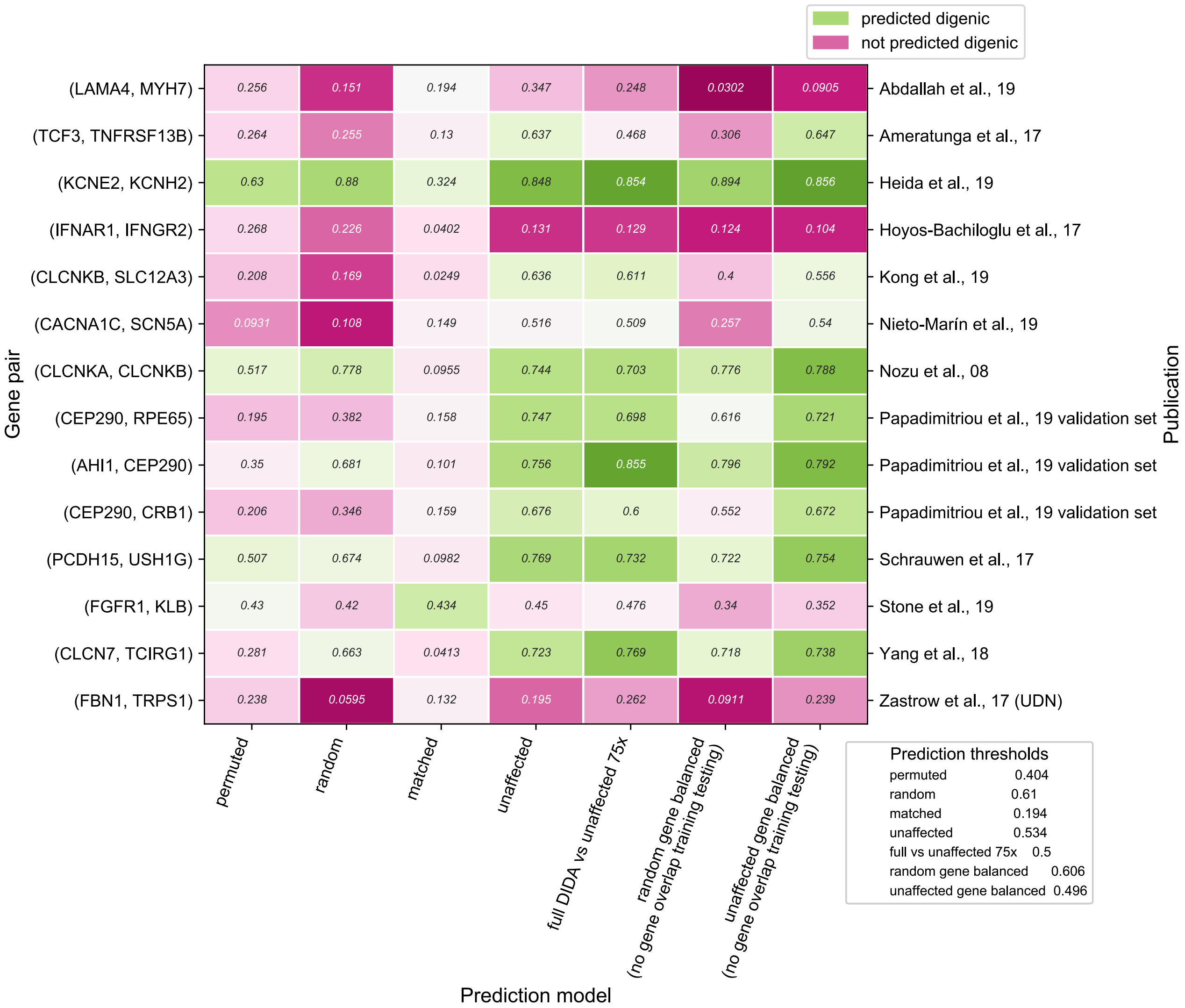


**FIGURE S9: VALIDATION OF OTHER MODELS OF CLASSIFIER USING NOVEL DIGENIC PAIRS FROM RECENT LITERATURE.**

The novel digenic pairs from recent literature and name of first author of the publication are along the Y axis. The predicted scores by the different models of DiGePred, trained on different negative sets, are along the X axis. Green indicates prediction as digenic based on $F_{0.5}$ prediction threshold, pink indicates no digenic prediction. 9/13 novel digenic pairs are identified as digenic by the unaffected model. Solved UDN case with overlapping phenotypes not predicted as digenic.

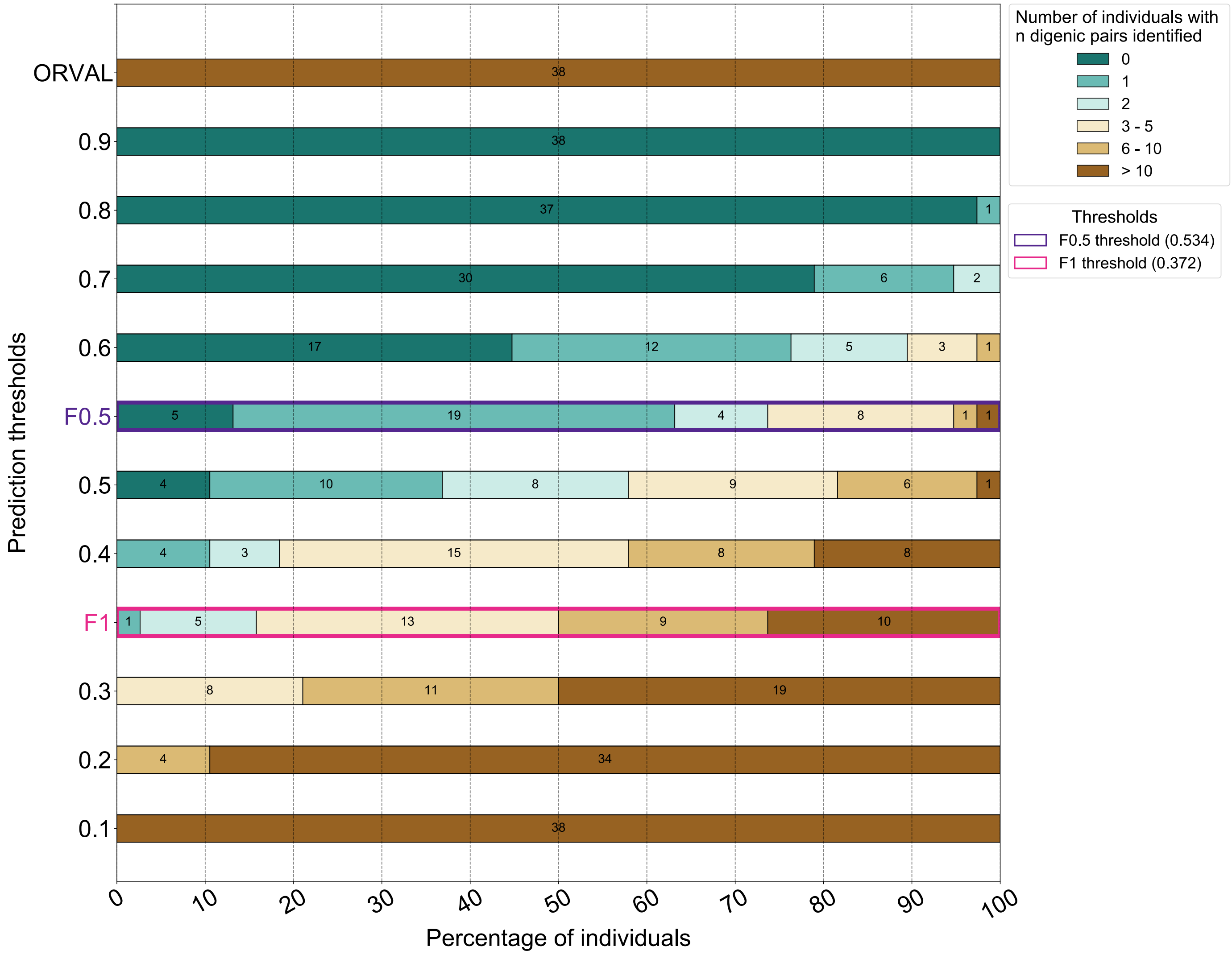


**FIGURE S10: LOW FALSE POSITIVE RATE OF CLASSIFIER ON EXTERNAL NEGATIVE TEST SET OF GENE PAIRS FROM UNAFFECTED RELATIVES OF UDN PATIENTS; COMPARISON WITH RECENTLY PUBLISHED VARIANT COMBINATION PATHOGENICITY PREDICTOR AT VARIOUS PREDICTION THRESHOLDS**

Number of gene pairs from individuals without digenic disease (unaffected relatives of UDN patients; n=38) identified to be digenic at varying predicted probability thresholds shown on X axis. Percentage of individuals with zero (dark green), one (green), two (light green), three - five (beige), six - ten (light brown) and > ten (dark brown) predicted digenic pairs shown on X axis; prediction thresholds shown on Y axis. ORVAL (VarCOPP) is a recently published variant combination pathogenicity predictor. The number in the box indicates number of individuals in each category. The prediction thresholds ($F_1$ score, shown in pink and $F_{0.5}$ score threshold shown in purple. At the $F_{0.5}$ threshold, five (13.2%) of unaffected individuals had no predicted digenic pairs, while 19 (50%) had only one predicted digenic pair. Only two individuals had more than five digenic pairs and only two digenic pairs were predicted per individual on average. All individuals were predicted to have more than ten digenic pairs by ORVAL, with an average number of 855 predicted digenic pairs per individual.

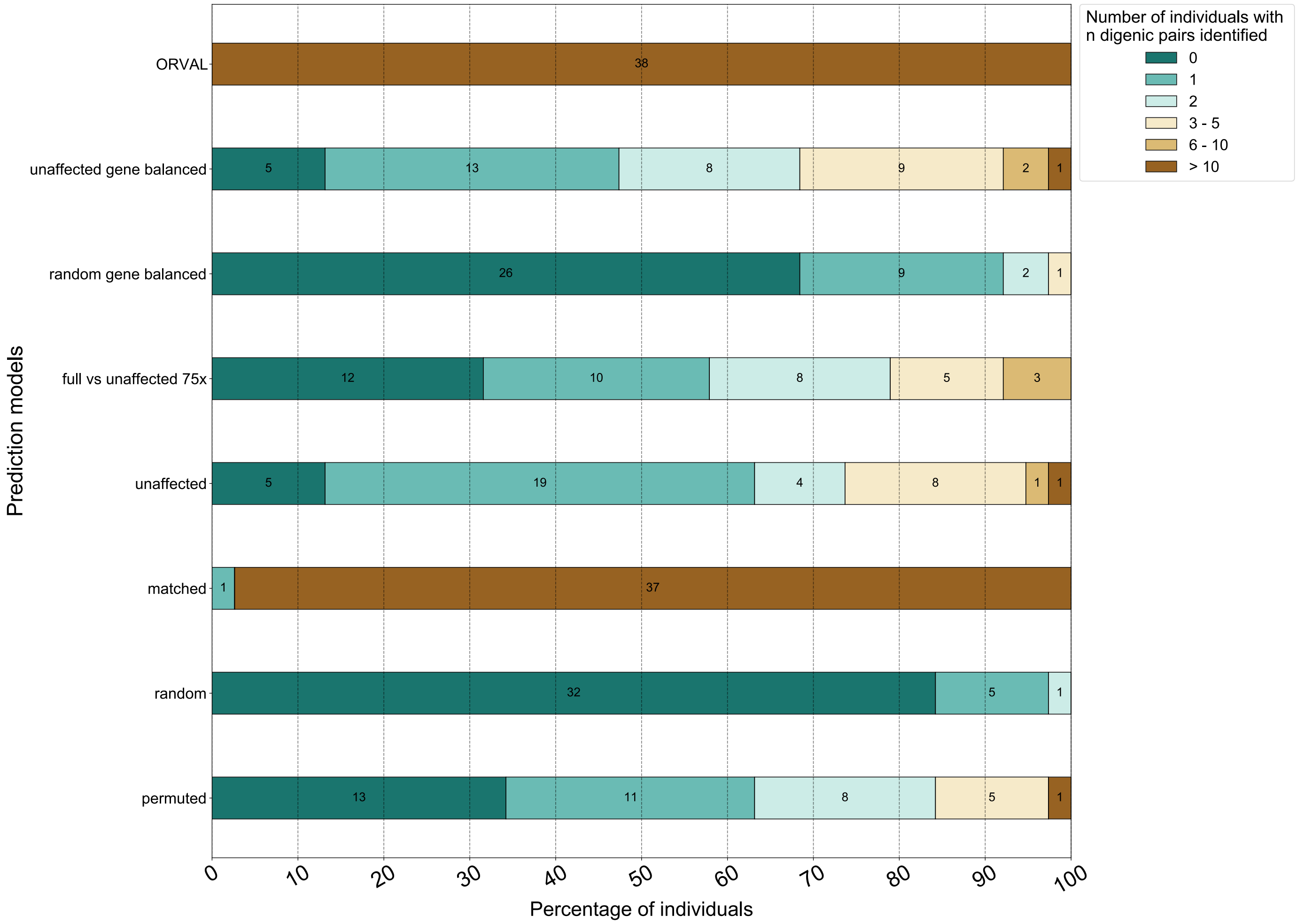


**FIGURE S11: LOW FALSE POSITIVE RATE OF CLASSIFIER ON EXTERNAL NEGATIVE TEST SET OF GENE PAIRS FROM UNAFFECTED RELATIVES OF UDN PATIENTS; COMPARISON WITH RECENTLY PUBLISHED VARIANT COMBINTAION PATHOGENICITY PREDICTOR USING VARIOUS MODELS OF TRAINING**

Number of gene pairs from individuals without digenic disease (unaffected relatives of UDN patients; n=38) identified to be digenic at varying predicted probability thresholds shown on X axis. Percentage of individuals with zero (dark green), one (green), two (light green), three - five (beige), six - ten (light brown) and > ten (dark brown) predicted digenic pairs shown on X axis; models of training on Y axis ORVAL (VarCOPP) is a recently published variant combination pathogenicity predictor. The number in the box indicates number of individuals in each category. At the $F_{0.5}$ threshold, all models except matched has fewer than three digenic pairs predicted for > 68% of unaffected individuals. ORVAL predicts more than ten digenic pairs for every individual.

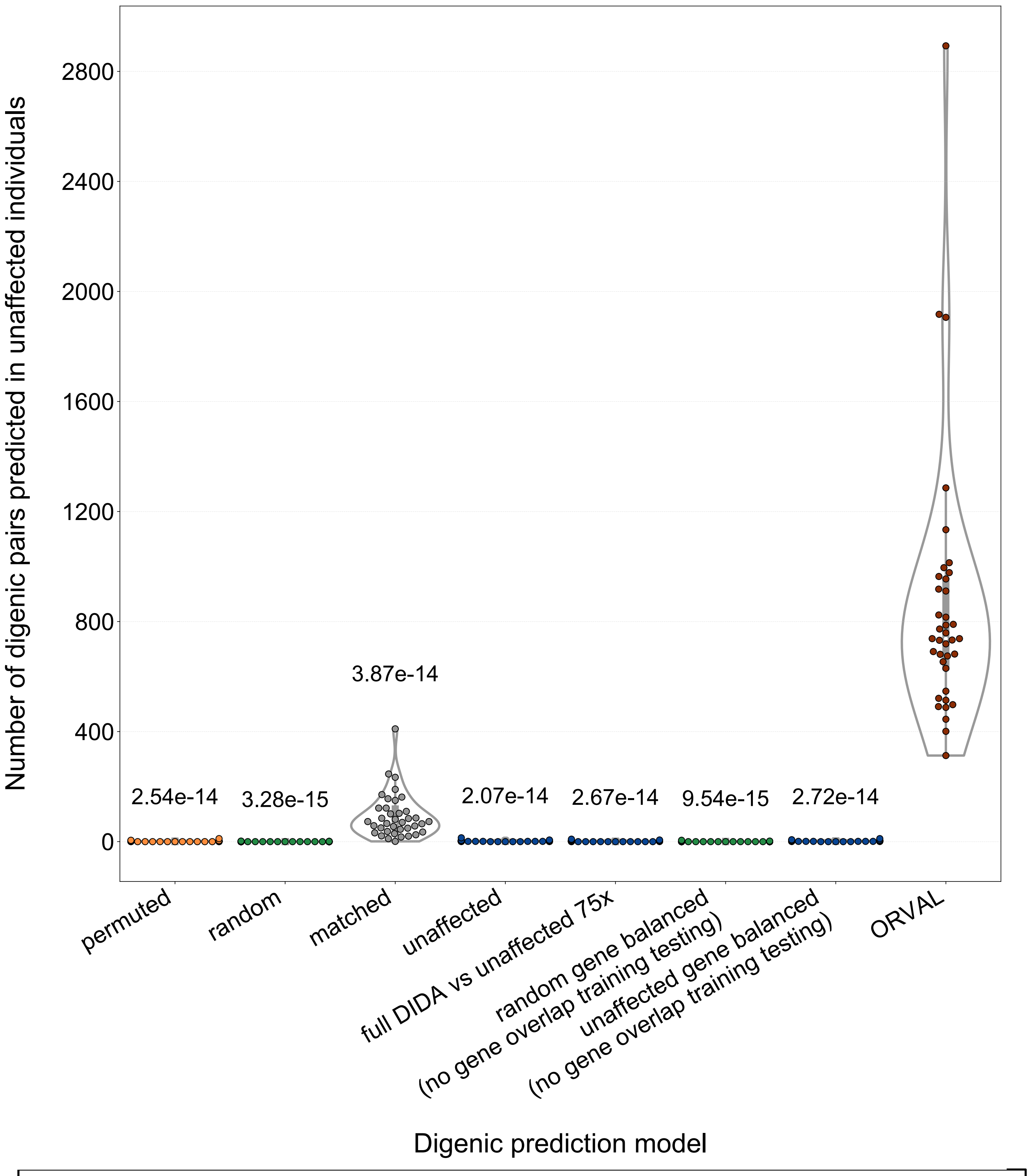


**FIGURE S12: FEWER FALSE POSITIVES FOR DiGePred COMPARED TO ORVAL FOR OTHER MODELS OF CLASSIFIER.**

The number of digenic pairs identified for each of 38 healthy relatives of UDN patients is plotted for different models of DiGePred, trained on different negative sets, (x-axis) and for the ORVAL/VarCOPP method. Since these individuals are healthy, any predicted digenic disease pairs a very likely false positives. DiGePred predicts significantly fewer digenic pairs for every model than ORVAL (MWU test, p-values above each bar). DiGePred trained on unaffected pairs predicts an average of two digenic pairs per healthy individual, while the permuted, random, matched, full DIDA, random gene balanced and unaffected gene balanced models predict an average of 1.5, 0.18, 92, 1.7, 1.47, 2.3 digenic pairs per individual respectively. ORVAL predicts an average of 855 digenic pairs per healthy individual.

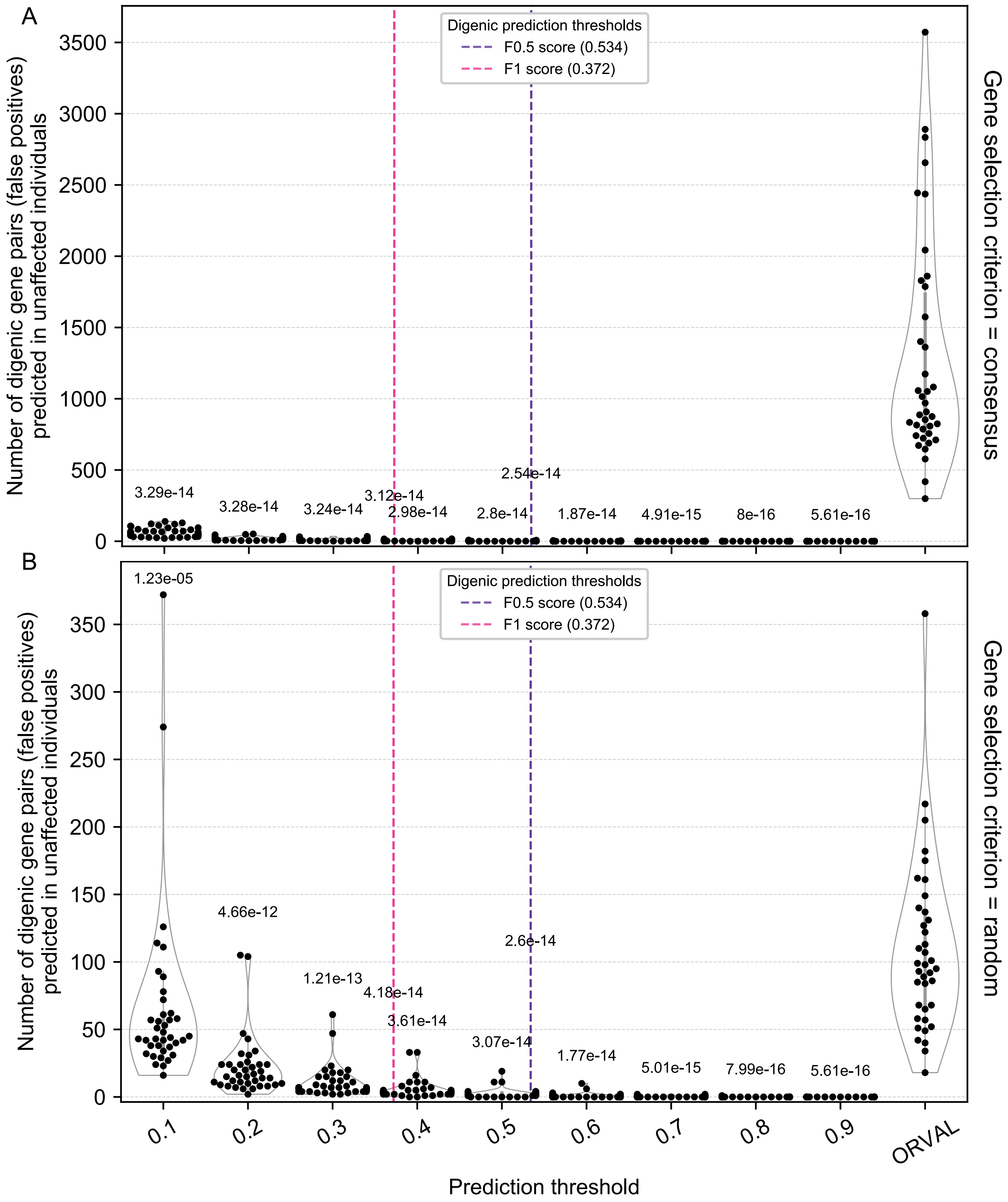


**FIGURE S13: FEWER FALSE POSITIVES FOR DiGePred COMAPRED TO ORVAL FOR OTHER GENE SECTION CRITERIA.**

The number of digenic pairs identified for each of 38 healthy relatives of UDN patients is plotted at a range of DiGePred thresholds (x-axis) and for the ORVAL/VarCOPP method. The score thresholds that maximize the $F_1$ and $F_{0.5}$ metrics on the held out data are shown in pink and purple, respectively. DiGePred predicts significantly fewer digenic pairs at each threshold than ORVAL (MWU test, p-values above each bar). The genes are selected by a **C**onsensus pathogenic criterion (**A**) and by **R**andom selection (**B**). At the F0.5 threshold, DiGePred predicts an average of 1.5(C) and 1.8 (R) digenic pairs per healthy individual, while ORVAL predicts an average of 1286 (C) and 108 (R) digenic pairs per healthy individual.

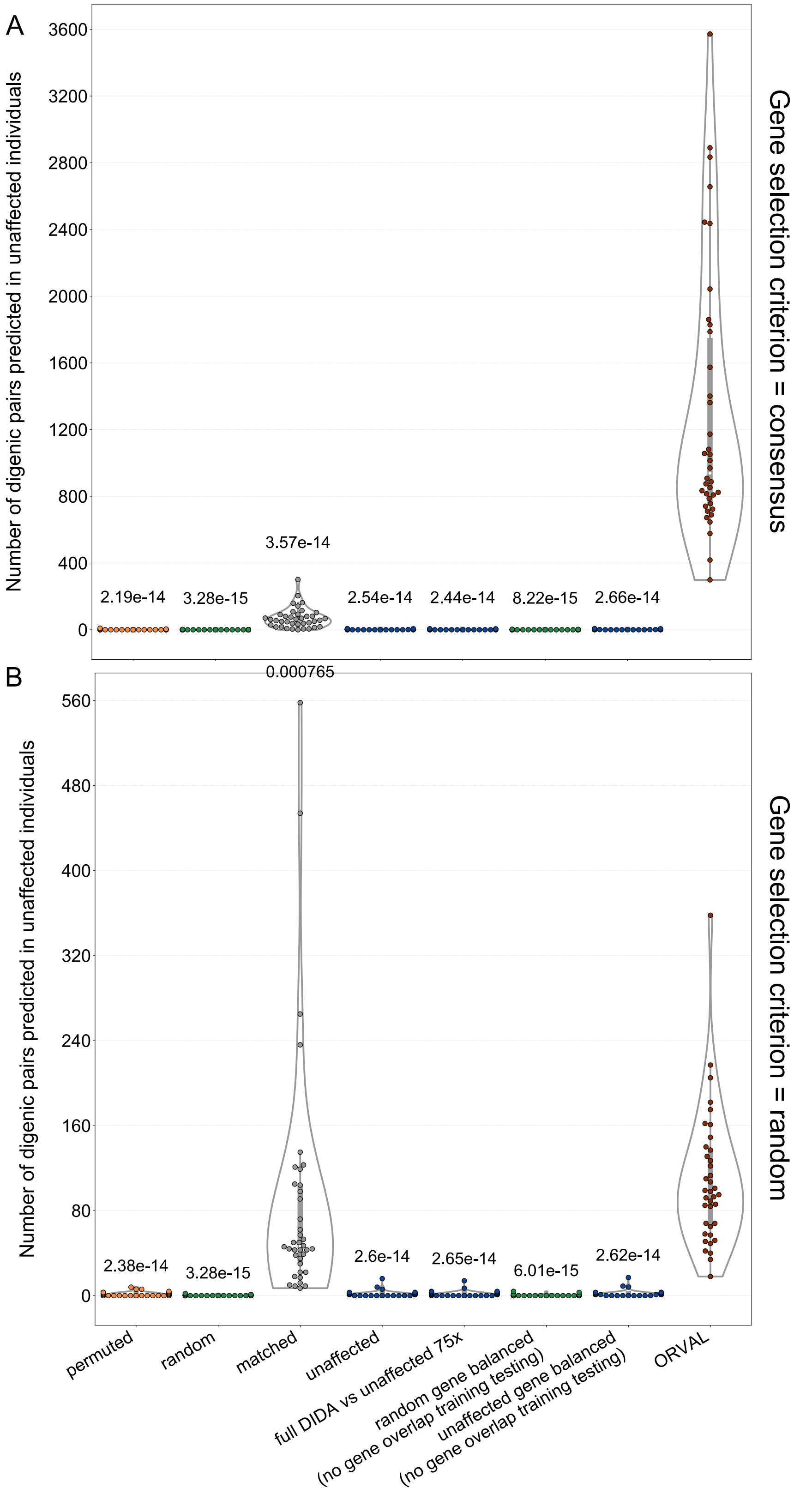


**FIGURE S14: FEWER FALSE POSITIVES FOR DiGePred COMPARED TO ORVAL FOR OTHER MODELS OF CLASSIFIER.**

The number of digenic pairs identified for each of 38 healthy relatives of UDN patients is plotted for **U**naffected, **P**ermuted, **R**andom, **M**atched, **F**ull DIDA, **R**andom gene **B**alanced and **U**naffected gene **B**alanced models of DiGePred, trained on different negative sets, (x-axis) and for the ORVAL method. DiGePred predicts significantly fewer digenic pairs for every model than ORVAL (MWU test, p-values above each bar). The genes are selected by a **C**onsensus pathogenic criterion (**A**) and by **R**andom selection (**B**). DiGePred predicts an average of 1.5C/1.8R (U), 1.1C/1.6R (P), 0.19C/0.19R (R), 70C/88R (M), 1.3C/1.7R (F), 0.37C/0.37R (RB) and 1.7C/2R (UB). ORVAL predicts an average of 1285C/ 108R.

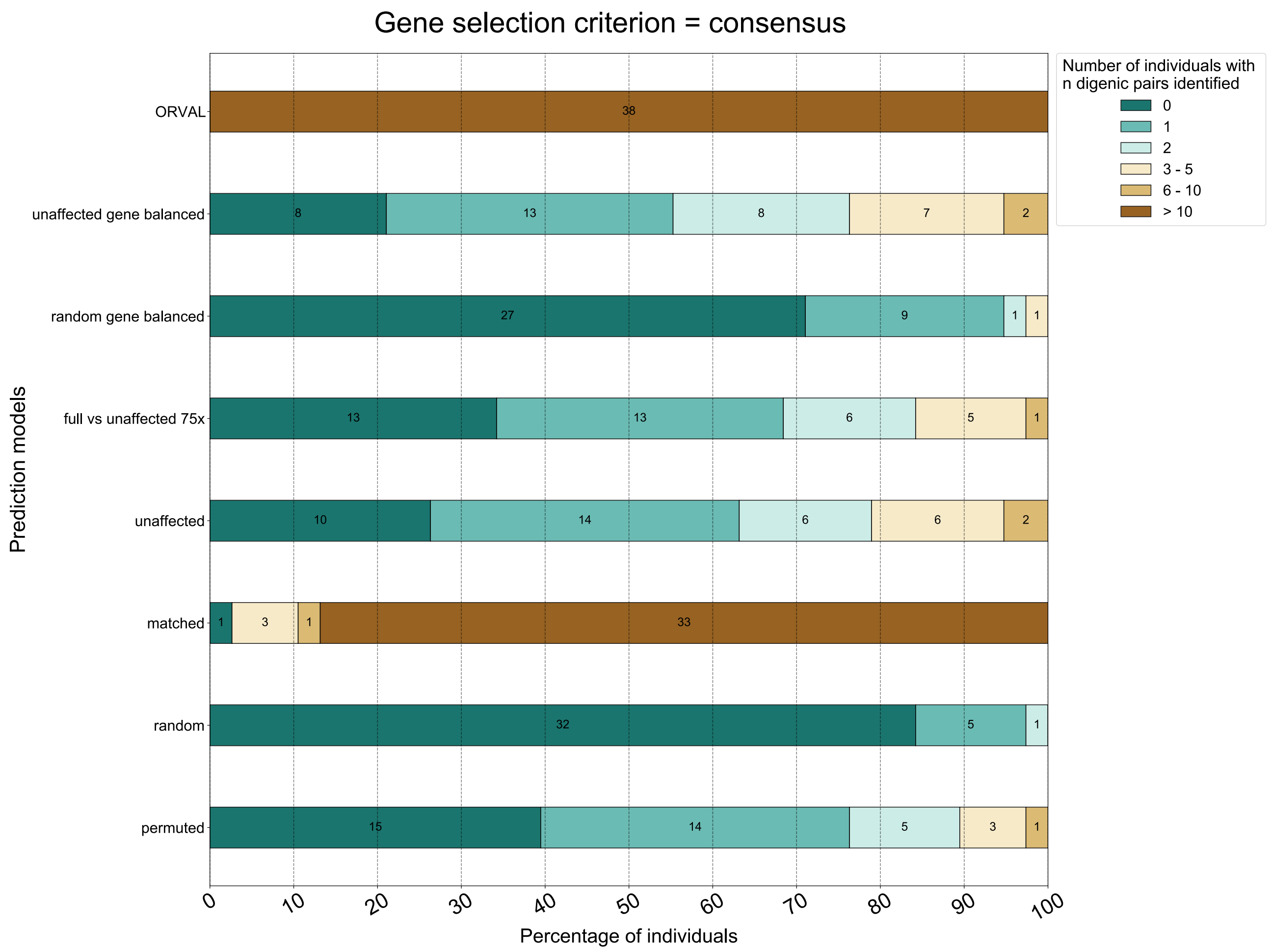


**FIGURE S15: LOW FALSE POSITIVE RATE OF CLASSIFIER ON EXTERNAL NEGATIVE TEST SET OF GENE PAIRS FROM UNAFFECTED RELATIVES OF UDN PATIENTS; COMPARISON WITH RECENTLY PUBLISHED VARIANT COMBINATION PATHOGENICITY PREDICTOR USING VARIOUS MODELS OF TRAINING**

Number of gene pairs from individuals without digenic disease (unaffected relatives of UDN patients; n=38) identified to be digenic at varying predicted probability thresholds shown on X axis. Percentage of individuals with zero (dark green), one (green), two (light green), three - five (beige), six - ten (light brown) and > ten (dark brown) predicted digenic pairs shown on X axis; models of training on Y axis ORVAL (VarCOPP) is a recently published variant combination pathogenicity predictor. The number in the box indicates number of individuals in each category. At the $F_{0.5}$ threshold, all models except matched has fewer than three digenic pairs predicted for > 76% of unaffected individuals. ORVAL predicts more than ten digenic pairs for every individual.

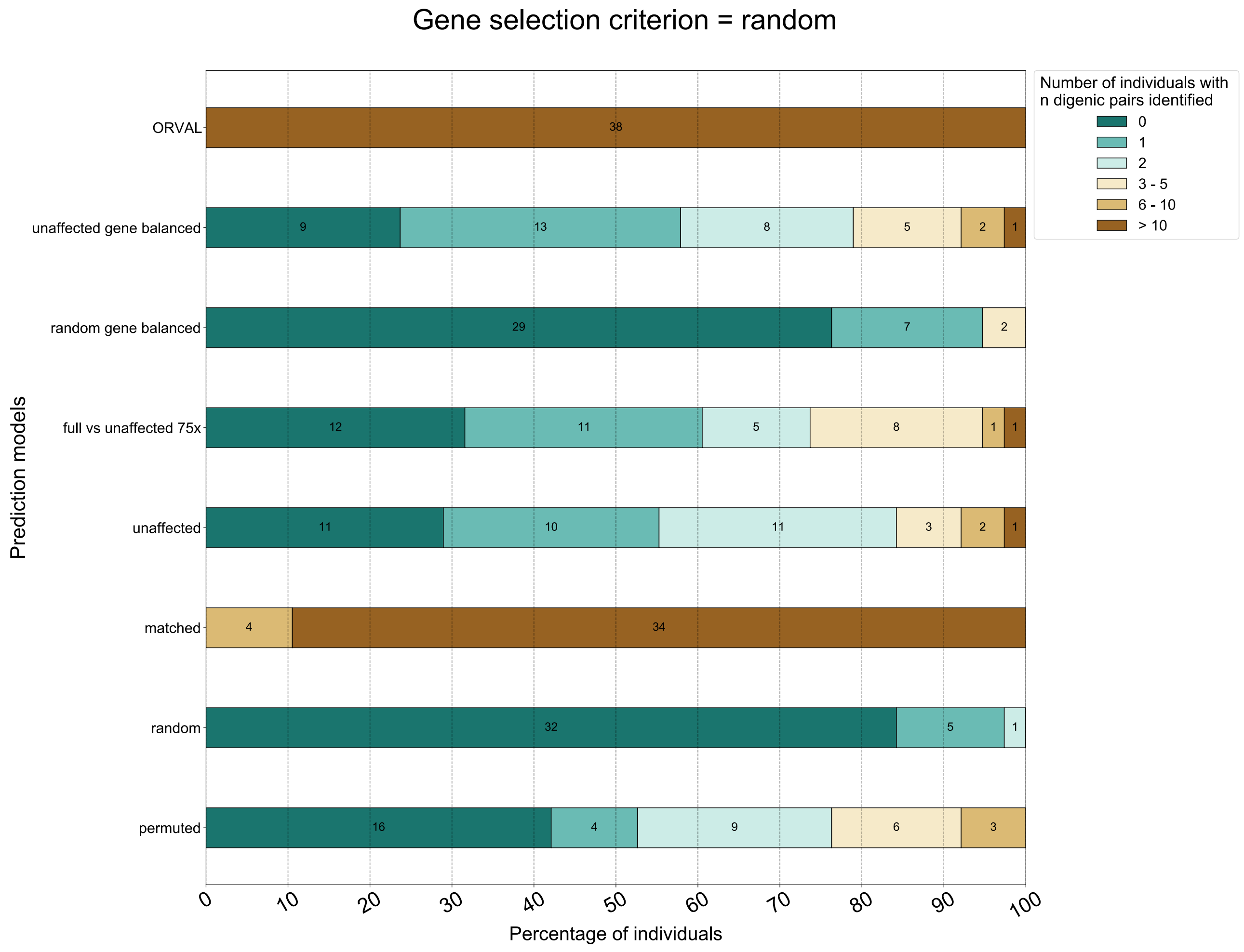


**FIGURE S16: LOW FALSE POSITIVE RATE OF CLASSIFIER ON EXTERNAL NEGATIVE TEST SET OF GENE PAIRS FROM UNAFFECTED RELATIVES OF UDN PATIENTS; COMPARISON WITH RECENTLY PUBLISHED VARIANT COMBINATION PATHOGENICITY PREDICTOR USING VARIOUS MODELS OF TRAINING**

Number of gene pairs from individuals without digenic disease (unaffected relatives of UDN patients; n=38) identified to be digenic at varying predicted probability thresholds shown on X axis. Percentage of individuals with zero (dark green), one (green), two (light green), three - five (beige), six - ten (light brown) and > ten (dark brown) predicted digenic pairs shown on X axis; models of training on Y axis ORVAL (VarCOPP) is a recently published variant combination pathogenicity predictor. The number in the box indicates number of individuals in each category. At the $F_{0.5}$ threshold, all models except matched has fewer than three digenic pairs predicted for > 73% of unaffected individuals. ORVAL predicts more than ten digenic pairs for every individual.

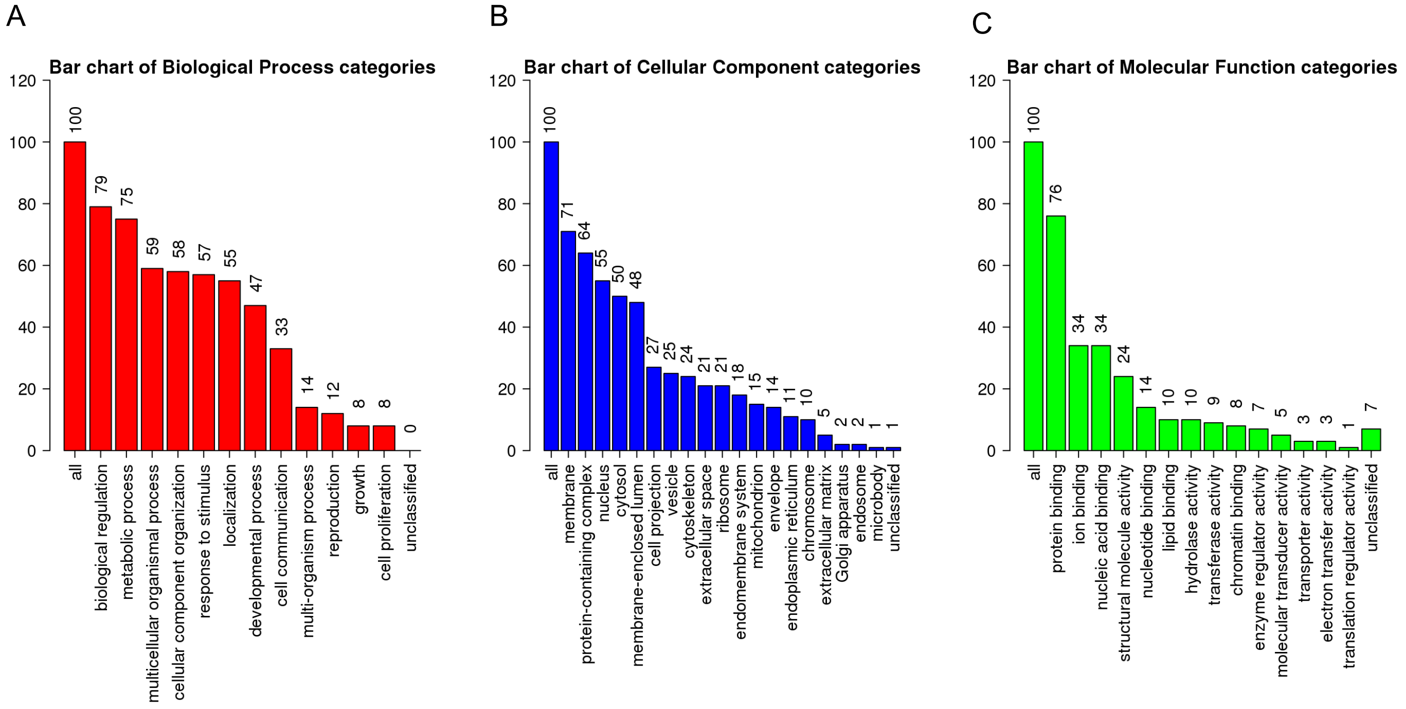


**FIGURE S17: GENE ONTOLOGY ENRICHMENT FOR TOP 100 GENES WITH MOST PREDICTED DIGENIC PAIRS**

Gene ontology (GO) enrichment using WebGestalt (WEB-based GEne SeT AnaLysis Toolkit). GO terms along X axis, and number along Y axis. (A) Biological process (red); (B) Cellular Component (blue); (C) Molecular function (green). Most genes are involved in metabolic processes, localized to the cell membranes and multi-protein complexes and have important binding domains.

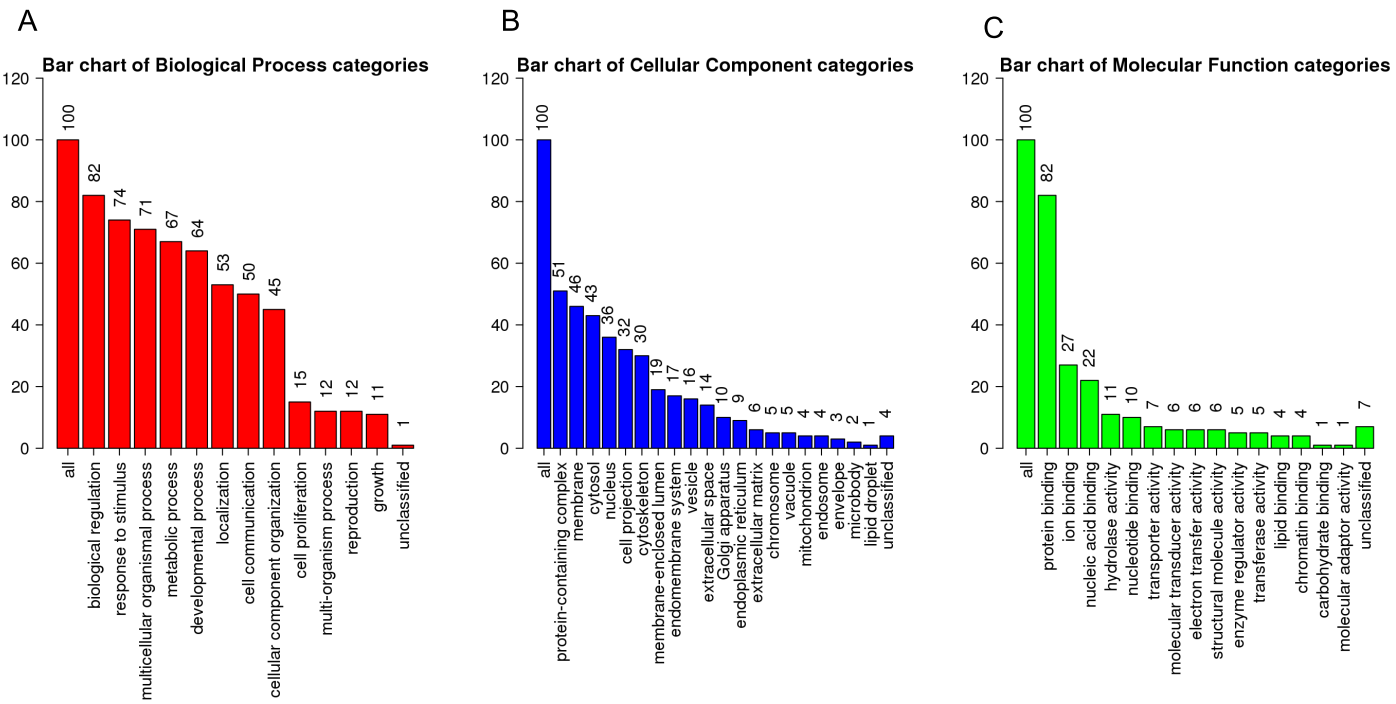


**FIGURE S18: GENE ONTOLOGY ENRICHMENT FOR TOP 100 GENES WITH HIGHEST AVERAGE PREDICTED VALUE**

Gene ontology (GO) enrichment using WebGestalt (WEB-based GEne SeT AnaLysis Toolkit). GO terms along X axis, and number along Y axis. (A) Biological process (red); (B) Cellular Component (blue); (C) Molecular function (green). Most genes are involved in metabolic processes, localized to the cell membranes and multi-protein complexes and have important binding domains.

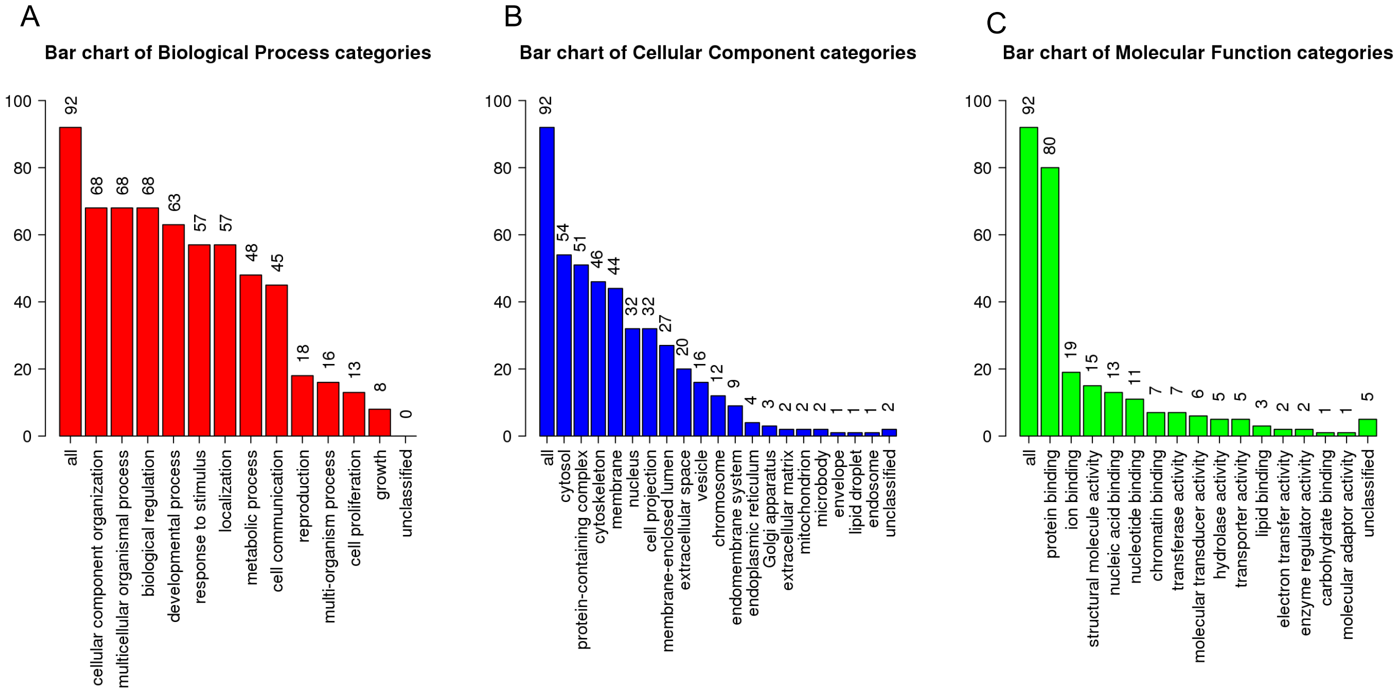


**FIGURE S19: GENE ONTOLOGY ENRICHMENT FOR GENES IN THE TOP 100 GENE PAIRS WITH HIGHEST PREDICTED VALUE**

Gene ontology (GO) enrichment using WebGestalt (WEB-based GEne SeT AnaLysis Toolkit). GO terms along X axis, and number along Y axis. (A) Biological process (red); (B) Cellular Component (blue); (C) Molecular function (green). Most genes are involved in metabolic processes, localized to the cell membranes and multi-protein complexes and have important binding domains.
